# Effect of Human Infant Gut Microbiota on Mouse Behavior, Dendritic Complexity, and Myelination

**DOI:** 10.1101/2023.10.24.563309

**Authors:** Harikesh Dubey, Rohon Roychoudhury, Ann Alex, Charlotte Best, Sheng Liu, Antonio White, Alexander Carlson, M. Andrea Azcarate-Peril, Linda S. Mansfield, Rebecca Knickmeyer

**Affiliations:** Institute for Quantitative Health Science and Engineering, Michigan State University, 775 Woodlot Drive, East Lansing, MI 48824; Department of Pediatrics, School of Medicine, University of California San Diego, 500 Gilman Drive, La Jolla, CA 92093; Department of Medicine, University of North Carolina at Chapel Hill, 125 MacNider Hall, Campus Box #7005, Chapel Hill, NC 27599; Department of Nutrition, Gillings School of Global Public Health, University of North Carolina at Chapel Hill, 135 Dauer Dr, Chapel Hill, NC 27514; Microbiome Core Facility, University of North Carolina at Chapel Hill, 312 Isaac Taylor Hall - Campus Box 7032, Chapel Hill, NC 27599; Departments of Large Animal Clinical Sciences and Microbiology and Molecular Genetics, Michigan State University, Food Safety and Toxicology Bldg., 1129 Farm Lane, East Lansing, MI 48824; Department of Pediatrics and Human Development, Michigan State University, Life Sciences Bldg. 1355 Bogue St., B240, East Lansing MI 48824

## Abstract

The mammalian gut microbiome influences numerous developmental processes. In human infants it has been linked with cognition, social skills, hormonal responses to stress, and brain connectivity. Yet, these associations are not necessarily causal. The present study tested whether two microbial stool communities, common in human infants, affected behavior, myelination, dendritic morphology, and spine density when used to colonize mouse models. Humanized animals were more like specific-pathogen free mice than germ-free mice for most phenotypes, although in males, both humanized groups were less social. Both humanized groups had thinner myelin sheaths in the hippocampus, than did germ-free animals. Humanized animals were similar to each other except for dendritic morphology and spine density where one group had greater dendritic length in the prefrontal cortex, greater dendritic volume in the nucleus accumbens, and greater spine density in both regions, compared to the other. Results add to a body of literature suggesting the gut microbiome impacts brain development.

**Teaser:** Fecal transplants from human infants with highly abundant *Bifidobacterium*, an important inhabitant of the intestinal tract of breastfed newborns, may promote brain connectivity in mice.

## Introduction

Environmental factors operating during the fetal and early postnatal period may contribute to future psychopathology by altering neurodevelopmental programming, in line with the Developmental Origins of Health and Disease (DOHaD) hypothesis (reviewed in (*1–3*)). While much work in this area has focused on prenatal stress and exposure to specific environmental chemicals, recent research suggests that the gut microbiota also influence brain development and subsequent motor, social, and cognitive functions (reviewed in (*4*, *5*)).

Initial evidence suggesting that gut microbes are involved in early programming of brain circuits came from studies of germ-free rodents, which differ from specific pathogen-free animals in their hormonal responses to stress, anxiety-like behavior, social interaction, and cognitive function (*6*–*10*). Such animals also display altered neurodevelopmental processes including neurogenesis, axonal and dendritic growth, and myelination (*6–11*). This suggests that gut microbes play a role in vulnerability to neuropsychiatric and neurodevelopmental disorders like schizophrenia and autism, where these processes and domains are often disrupted. This hypothesis is further supported by numerous studies showing significant differences in individuals’ microbial communities with and without these conditions (*12–17*).

Infancy is an important period in the assembly and maturation of the gut microbiome. At birth, the human infants born vaginally have guts with a low abundance lowly diverse microbial community reflective of the mother’s vaginal and fecal microbiota, then progresses through three phases: a developmental phase (months 0-14), a transitional phase (months 15-30), and a stable phase (30 months and beyond) (*18*). Mother-to-infant microbial transmission plays a key role in this process with the maternal gut microbiome being a key donor of infant-acquired strains (*19*, *20*). The developmental phase of the gut microbiome coincides with the most rapid and dynamic phase of postnatal brain development (*21*), suggesting variation in this period could have long-term impacts on brain structure and function (*22*). Indeed, our group has demonstrated that certain features of the infant gut microbiome are associated with cognition (*23*), non-social fear reactivity behavior (*24*), hormonal responses to stress (*25*), regional brain volumes (*23*, *24*), and functional brain connectivity (*28*). Given the observational nature of our human infant studies, a causal relationship between the human infant gut microbiome and individual differences in neurodevelopment and behavior has yet to be established. To better understand how different human infant gut microbiomes influence neurodevelopment, we carried out the present study to investigate whether transplantation of human infant stool communities into germ-free pregnant mice altered mouse behavior and dendritic complexity and myelination in brain regions implicated in emotional reactivity and social behavior including the amygdala, hippocampus, nucleus accumbens, and prefrontal cortex. We focused on two communities commonly found in human infants’ stool: one with a relatively high abundance of *Bifidobacterium* (HUM1) and one with a relatively high abundance of *Bacteroides* (HUM2). Microbial communities were introduced into pregnant germ-free animals at mid-gestation (pregnancy day 10) to ensure the offspring were exposed from birth onwards. For comparison purposes, we also inoculated a group of pregnant germ-free animals with an inoculum created from the feces of specific-pathogen free mice (SPF) and another group with autoclaved fecal material from SPF mice. The latter group remained germ-free (GF) until the time of behavioral testing. Three different cohorts were generated for approximately one year to ensure sufficient animals were studied. Engraftment was determined by 16S rRNA gene sequencing analysis.

## Results

### Microbial taxonomic composition differed between humanized mice and specific-pathogen free and germ-free controls

16S rRNA amplicon sequencing was used to compare microbial taxonomic composition between groups using cecal contents collected post-mortem (around 9 weeks of age). Alpha (within sample) diversity was assessed using the Shannon index and Chao1 and differed across microbiome inoculation groups (Suppl Data S1 (a-b)). The Shannon index, which measures the evenness of species in a community, was significantly lower in humanized mice compared with SPF mice. Chao1, a nonparametric estimator of species richness, was significantly lower in HUM2 animals as compared to SPF animals. This mirrors differences in the original inocula as both HUM1 and HUM2 inocula had lower alpha diversity than the mouse SPF inoculum across various measures (Suppl S1(c)). Beta diversity was visualized using PCoA based on unweighted and weighted UniFrac, Bray-Curtis, and Jaccard distance matrices and suggested variation between individuals was influenced both by microbiome inoculation group (HUM1, HUM2, SPF, GF) and the date on which samples were collected (corresponds with cohort). PERMANOVA confirmed significant differences in microbial community between inoculation groups and between cohorts [Suppl data S1 (d-g)]. Similar patterns were observed in analyses stratified by sex. Thus, cohort was included as a covariate in the behavioral analyses and analyses of dendritic and myelination phenotypes.

We then performed linear discriminant effect size analysis (LEfSe) to identify which specific bacteria predicted membership in each microbiome inoculation group. *Bifidobacterium* and *Turicibacter* predicted inclusion in the HUM1 group. *Bacteroides* and Lachnospiraceae_NK4A136 predicted inclusion in the HUM2 group. *Lactobacillus* and *Alistipes* predicted inclusion in the SPF group, and *Faecalibaculum* and *Lachnospiraceae* predicted inclusion in the GF group of animals (Figure 1a,b). Variation in the relative abundance of microbial genera per sample was visualized using bar plots (Suppl S1(h and i)). Further, we examined shared operational taxonomic units (OTUs) across different groups (Figure 1c). The GF group had the largest number of unique OTUs (8%), and the SPF group had the lowest (6%), while 31% of OTUs were shared among all the groups.

**Figure 1:**
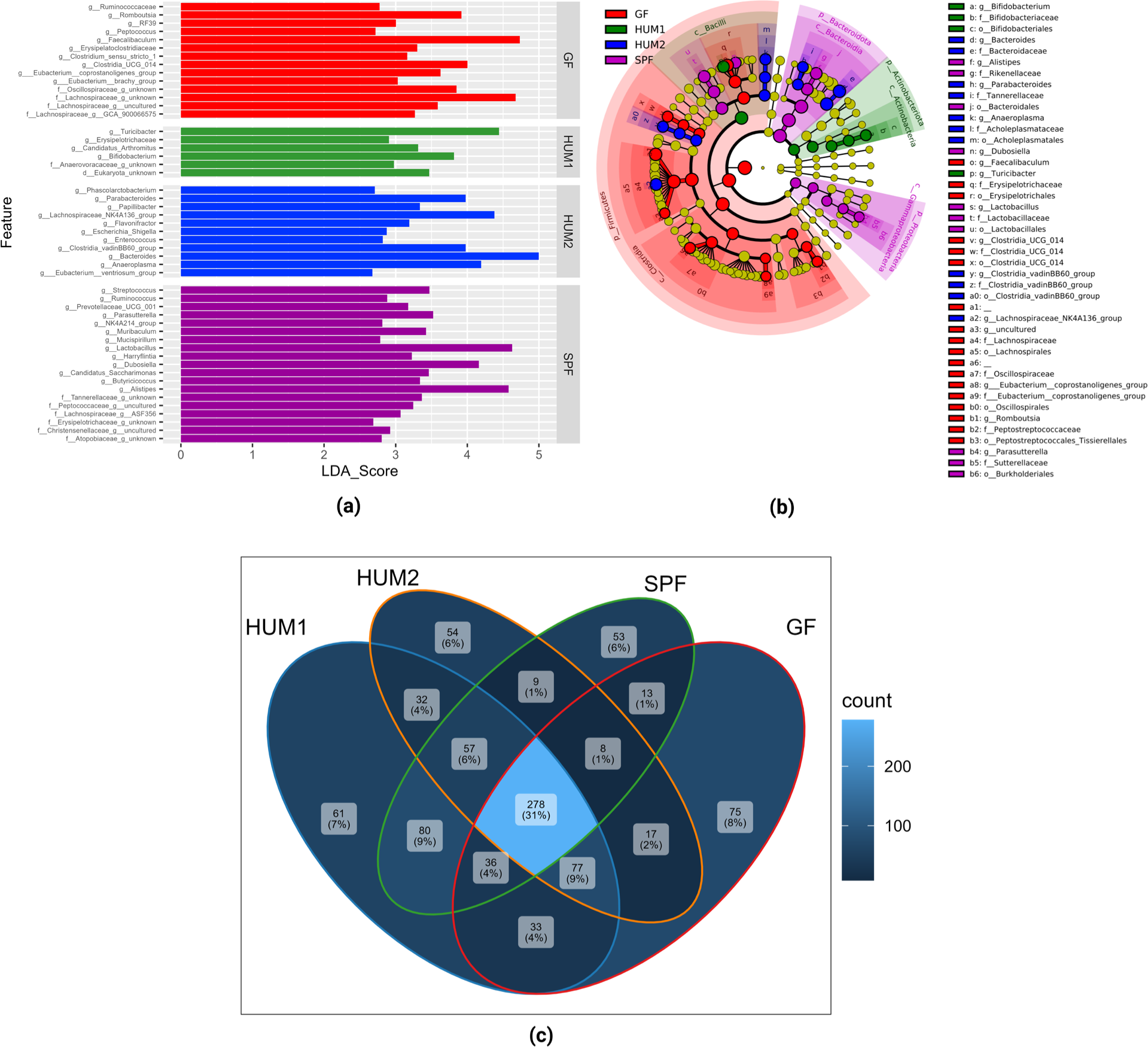
Microbial taxonomic composition differs at the genus level between GF, HUM1, HUM2, and SPF groups. LDA effect size (LEfSe) score (a), LEfSe cladogram (b) and Venn diagram, illustrated observed overlap of OTUs of the gut microbiota in GF, HUM1, HUM2, and SPF groups (c). Created with BioRender.com

To further determine how successfully infant gut microbiomes colonized the relevant mice groups, as per Aluthge et al. (2020), we assessed how many amplicon sequence variants (ASVs) present in the original inocula were detected in the cecal contents of the corresponding group. If a donor ASV was detected in more than 50% of recipient animals’ offspring, then this ASV was considered a “persistent colonizer” (*26*). Around 60% of ASVs in the SPF inoculum were persistent colonizers, while around 40% of ASVs in HUM1 and HUM2 were persistent colonizers (Supp data S1(j)).

Overall, microbiome patterns observed in the original inocula were partially, but not completely replicated in the mice, which is similar to prior studies using human microbiota-associated mouse models.

### Microbiome inoculation group influenced anxiety-related and exploratory/locomotor behaviors, but not social behavior, recognition memory, or depression-related behavior

Anxiety-related and exploratory/locomotor behaviors were evaluated using the elevated plus maze (EPM) (*27*, *28*), light/dark test (LDT) (*29*, *30*), and the open field test (OFT) (*31*, *32*). In the EPM, microbiome inoculation groups (HUM1, HUM2 and SPF) differed significantly (<0.0026) in time spent in the closed arm (F_3,38_ = 11.613, p = 0.0000152, three-way ANOVA) and number of entries into the closed arm F_3,38_ = 5.816, p = 0.00225, three-way ANOVA). Post-hoc pairwise comparisons indicated that GF mice spent less time in and made fewer entries into the closed arm than the other three groups (Figures 2a and 2b). A reduced preference for remaining in enclosed spaces is generally considered to indicate reduced anxiety. No significant differences were observed between microbiome inoculation groups in the number of entries in the open arm, time spent in the central arm, total time spent and total distance traveled in the open arm, or distance traveled in the closed arm and central arm (Suppl S2 (a)). Sex-stratified analyses indicate this pattern is present in both males and females for time spent in the closed arm (Figure 2c and 2d) and only in females for the number of entries (Figure 2e and 2f). In the LDT, microbiome inoculation groups differed significantly (<0.0026) in number of entries into the light chamber from the dark chamber (F_3,38_ = 6.256, p = 0.00147, three-way ANOVA). GF mice entered the dark chamber more often than the other groups, though this fell just short of significance for pairwise comparison with HUM2 animals (Fig 2g). Because no significant differences were observed for time spent in the dark chamber (an index of anxiety) (Suppl S2 (b)), the greater number of entries indicates greater locomotor activity/exploratory behavior in the GF mice. Sex-stratified analyses indicate this pattern is present in both males and females (Fig 2(h and i)). No significant group differences were observed for OFT phenotypes including time spent in the outer zone, distance traveled in the outer zone, velocity in the outer zone, time spent in the central zone, distance traveled in the central zone, velocity in the central zone, and total fecal boli count (Suppl S2 (c)).

**Figure 2:**
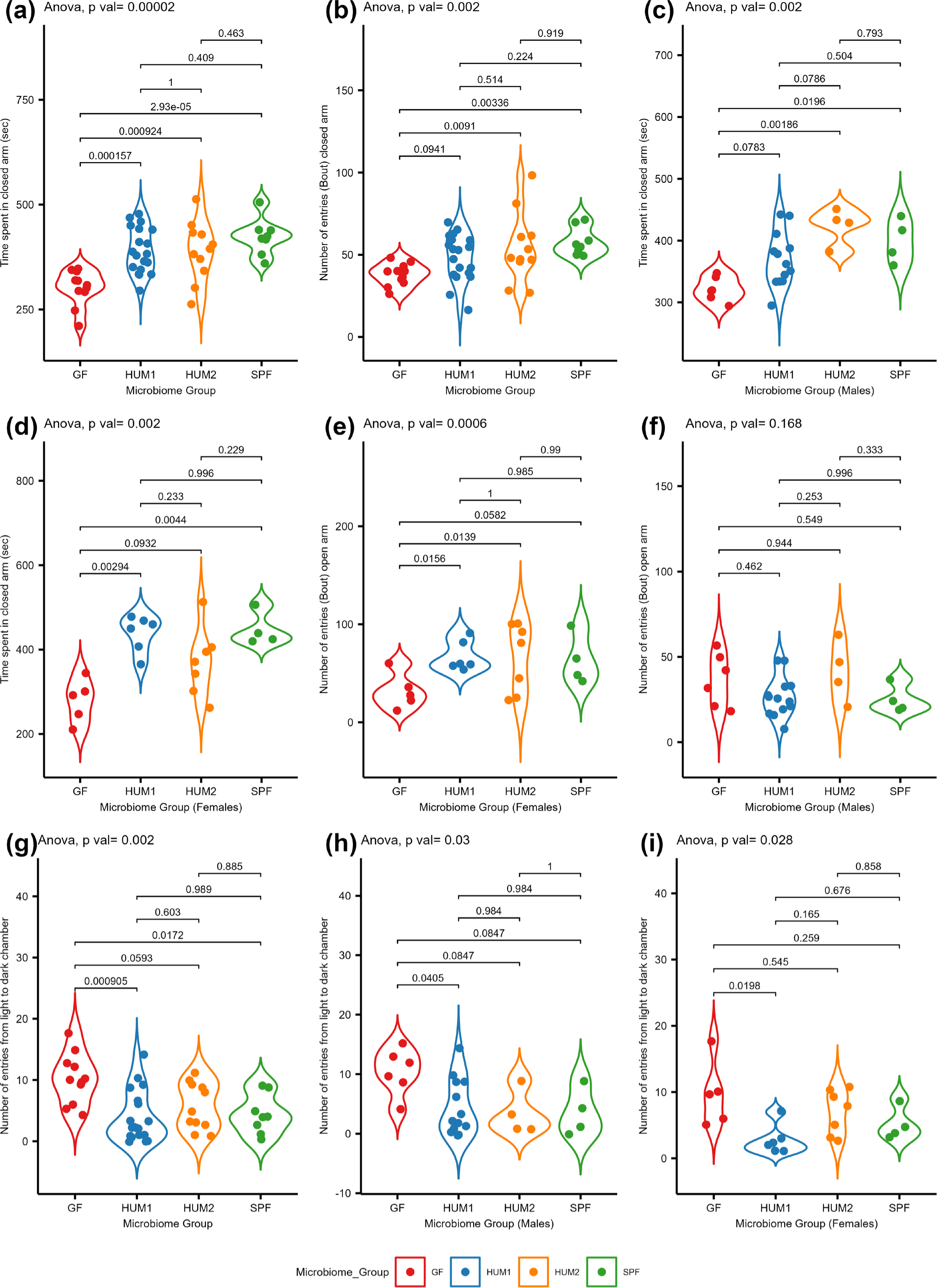
Effect of gut microbiomes on anxiety behavior. (a-e) Elevated plus maze behavioral tests, (a) time spent in the closed arm (M+F), (b) number of entries in the closed arm (M+F), (c) time spent in the closed arm (male), (d) time spent in the closed arm (female), (e) number of entries in open arm (female), (f) number of entries in open arm (male). (g-i) Light/dark preference test, (g) number of entries in from light chambers to dark chamber (M+F), (h) number of entries in from light chambers to dark chamber (male), (i) number of entries in from light chambers to dark chamber (female).

Recognition memory was evaluated using a novel object recognition (NOR) test (*33*). Microbiome inoculation group did not significantly impact outcomes on this test, which included time spent exploring the familiar object, time spent exploring the novel object, and the novel object discrimination index (Suppl S2 (d)). Sociability and preference for social novelty were investigated using the three-chamber social test (*34*). Our primary analyses (Three-way ANOVA) showed no significant impact of microbiome inoculation group on sociability as indexed by time spent exploring a novel con-specific in phase 1, when the novel mouse is in one chamber and an empty cage is in another chamber (Suppl S2 (e)). Similarly, there was no impact of the microbiome inoculation groups on preference for social novelty in phase 2 indexed by time spent exploring the familiar mouse, time spent exploring the novel mouse, and the novel mouse discrimination index. There was a strong effect of the cohort on time spent exploring the familiar mouse and time spent exploring the novel mouse in Phase 2 (Suppl S2 (e)), which may have limited our ability to observe the impact of the microbiome inoculation group. In addition, no group showed a strong preference for social novelty, which is unusual for this test. Although the three-way ANOVA did not indicate a significant interaction between the microbiome inoculation group and sex, sex-stratified analyses revealed that the microbiome inoculation group may influence male social behavior (F_3,22_ = 10.19, p = 0.000209, Two-way ANOVA). Male GF and SPF mice spent more time with the familiar mouse in Phase 2 than humanized mice (Fig 3). They also spent more time with the novel mice, though this was not significant (F_3,22_ = 2.66, p = 0.07332, Two-way ANOVA). Discrimination index was similar across groups suggesting male humanized mice have reduced sociability rather than a change in preference for social novelty.

**Figure 3:**
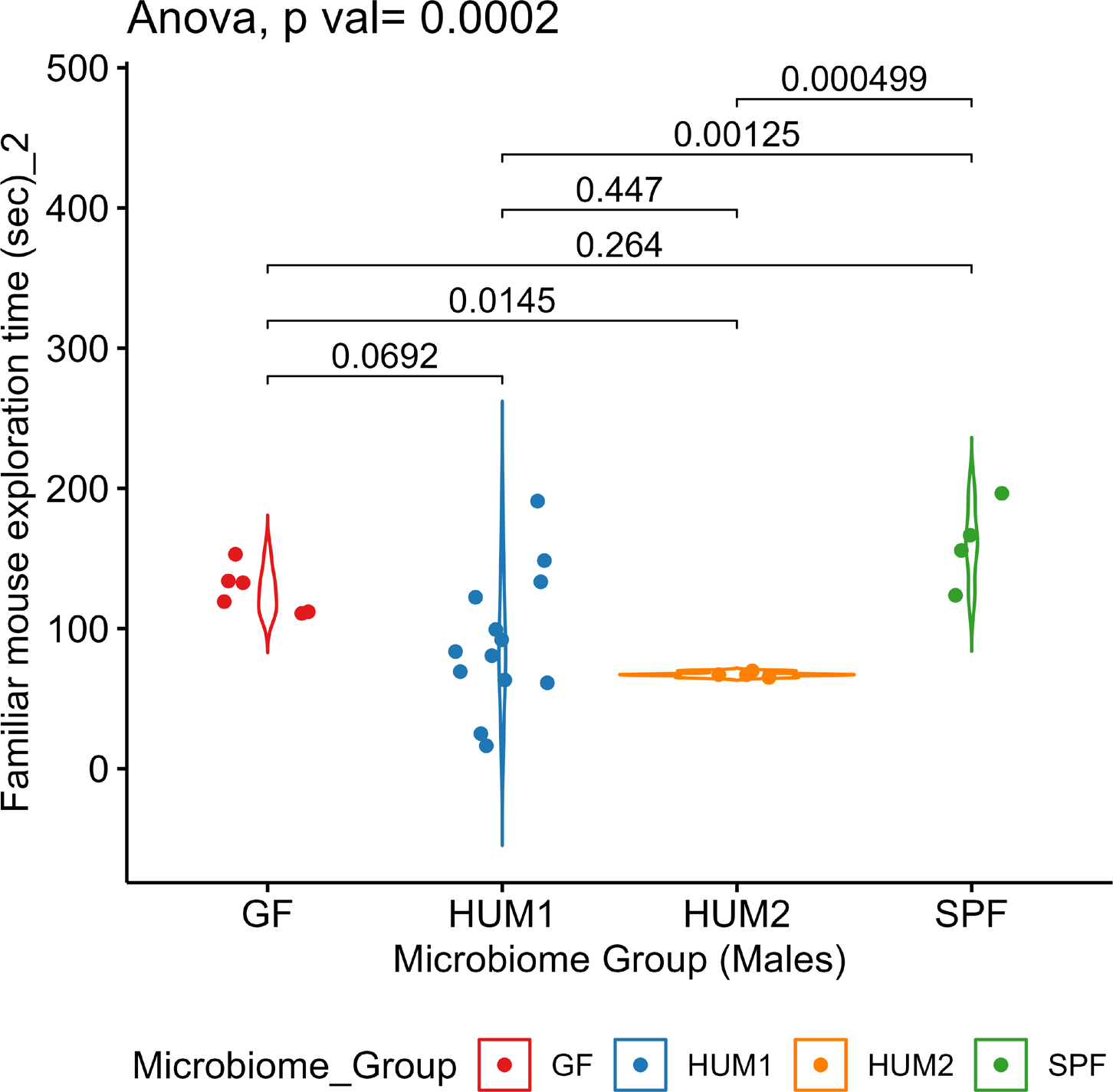
Three chamber sociability tests, familiar mouse exploration time (male).

Depression-related behavior was assessed with the sucrose preference test (SPT) (*35*, *36*). Microbiome inoculation group had a significant impact (p < 0.05) on sucrose preference (F_3,36_ = 4.083, p = 0.0136, mixed model ANOVA with repeated measures) as did day of consumption (F_2,72_ = 14.504, p = 0.000051, mixed model ANOVA with repeated measures). Sucrose preference increased over time, as is typical. While none of the post-hoc comparisons were significant, GF animals had lower preference for sucrose than SPF and HUM1 animals across all days of the test, with HUM2 being intermediate between those groups (Fig 4a). Moreover, similar results were observed in sex-stratified analyses of female mice (F_3,15_ = 3.74, p = 0.0347, mixed model ANOVA with repeated measures, but no significant variation was observed during post hoc-t test comparison) (Fig 4b). However, microbiome inoculation did not affect SPT in male mice (F_3,15_ = 2.094, p = 0.132, mixed model ANOVA with repeated measures), (Suppl S3 (a)).

**Figure 4:**
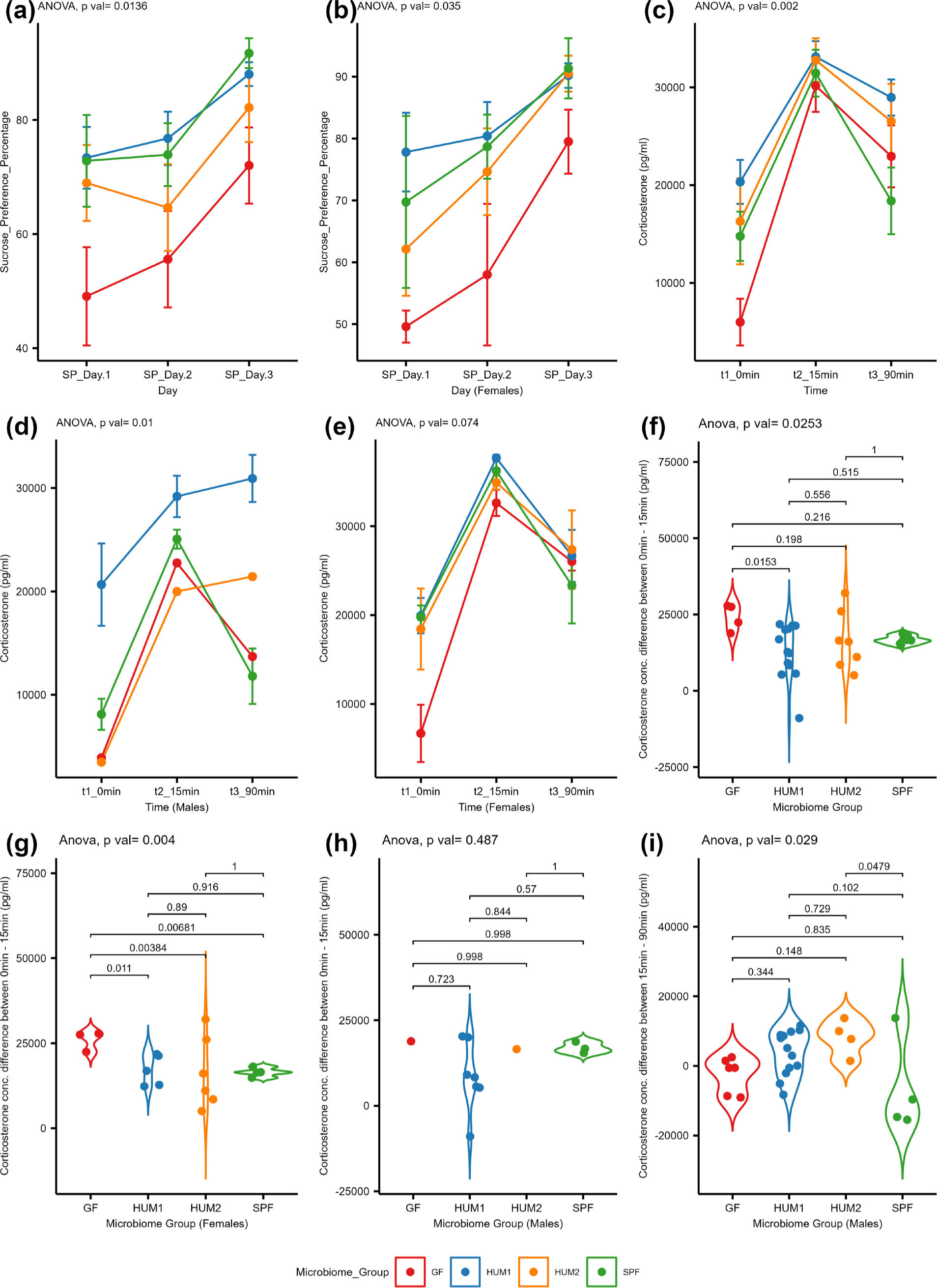
Effects of gut microbiome inoculation group on the sucrose preference test (a, b) and corticosterone levels during stress reactivity test (c-i), (a) Percentage sucrose preference intake by mice during day 1, 2 and 3 (cohort wise M+F), (b) Percentage sucrose preference intake by female mice during day 1, 2 and 3, (c) time dependent corticosterone levels-cohort wise (M+F) [significant level at ‘0’ min (GF vs HUM1 p=0.0377)], (d) time dependent corticosterone level variations in male mice (did not perform the posthoc test due to lack of sufficient number of samples), (e) time-dependent corticosterone level variations in female mice, (f) corticosterone level variations between 0-15min (combined gender, cohort wise), (g) corticosterone level variations between 0-15min in female mice, (h) corticosterone level variations between 0-15min (male), (i) corticosterone level variations between 15-90min (male).

Hormonal responsivity to stress was assessed with a restrain stress test (SRT) (*37*). A mixed model ANOVA with repeated measures revealed that microbiome inoculation group influenced stress reactivity (F_3,20_ = 7.358, p = 0.00164). Posthoc tests indicated that HUM1 animals had significantly higher corticosterone levels than GF animals at baseline (p=0.0377) (Fig 4c). HUM1 animals also had higher corticosterone levels than SPF at 90 minutes (recovery), though this did not reach statistical significance (p=0.0565). Although the three-way ANOVA did not indicate a significant interaction between microbiome inoculation group and sex, sex-stratified analyses suggest the impact of gut microbiome inoculation group on the SRT may be stronger in males (F_3,15_ = 8.33, p = 0.0104, mixed model ANOVA in males with repeated measures) versus females (F_3,15_ = 2.928, p = 0.0735) (Fig 4b and c). To better understand the significant microbiome effects on stress reactivity revealed in the mixed model repeated measures ANOVA, we conducted secondary analyses looking at microbiome effects on change in corticosterone from baseline to 15 minutes (animal still in restraint tube), change from 15 minutes to 90 minutes (animal no longer in tube), and change from baseline to 90 minutes. These analyses suggested that microbiome inoculation group influenced initial stress responses, indexed by change from baseline to 15 minutes (F_3,20_ = 3.843, p = 0.0253, three-way ANOVA), especially in females (F_3,13_ = 7.403, p = 0.00386, two-way ANOVA in females versus F_3,7_ = 0.901, p = 0.487 in males) (Fig 4f, g and h). Post hoc pairwise comparisons in the combined sex group show that GF mice have stronger increases in corticosterone from baseline to 15 minutes than HUM1 animals. Post hoc pairwise comparisons in females show that GF animals have stronger increases in corticosterone from baseline to 15 minutes than all other groups. However, during recovery (15 to 90 minutes) period, neither combined sex group (F_3,38_ = 2.521, p = 0.072362, three-way ANOVA) nor female mice (F_3,16_ = 0.298, p = 0.827, two-way ANOVA) showed significant variations in corticosterone levels by microbiome inoculation group (Suppl S3 (c)). Only male animals showed significant differences between microbiome inoculation groups in corticosterone change during recovery (F_3,22_ = 3.612, p = 0.0293, two-way ANOVA, Fig 4i). Post hoc comparisons indicated that male SPF animals recovered better than other groups with most SPF males showing a drop in corticosterone from 15 to 90 minutes. The difference between SPF males and HUM2 males was significant (p=0.0479) with corticosterone levels continuing to rise in all HUM2 animals.

### Microbiome inoculation group had minimal influence on gut function as indexed by intestinal transit time

Intestinal transit time was measured using the red carmine test (*38*) in which mice are gavaged with non-absorbable red carmine dye and the time to the first appearance of the dye in feces is recorded. Group differences in intestinal transit time were not significant (Suppl S4). The three-way ANOVA did not indicate a significant interaction between microbiome inoculation group and sex. However, in sex-stratified analyses the two-way ANOVA indicated the effects of the microbiome inoculation group in males (F_3,22_ = 3.658, p = 0.028). Posthoc pairwise comparisons were not significant. As a group, GF and SPF males may have longer transit times than humanized males, however this could reflect the influence of one GF male and one SPF male with longer transit times (Fig 5).

**Figure 5.**
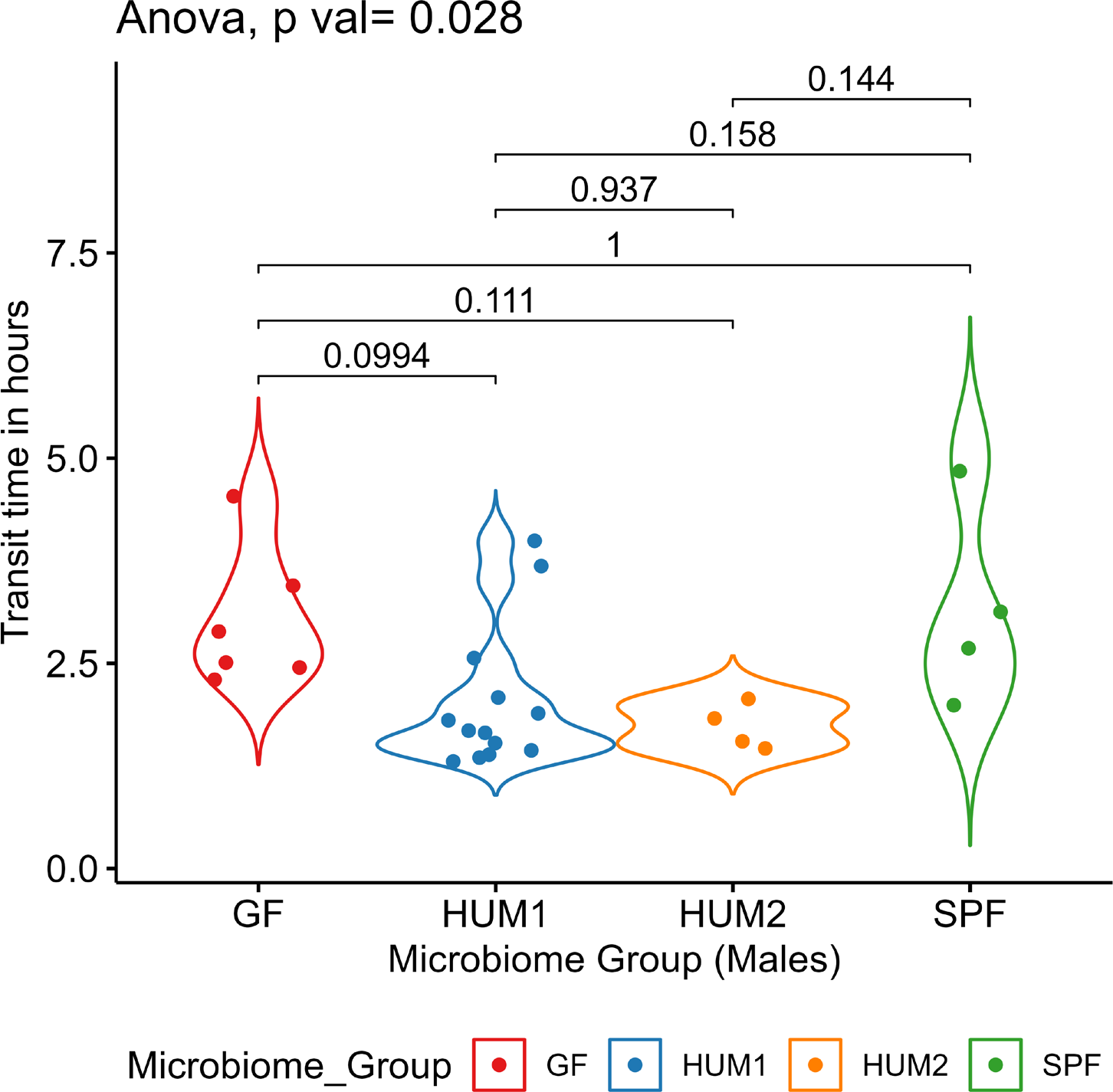
Effect of microbiome on intestinal transit time in mice (male).

### The gut microbiome may influence myelin sheath thickness in the hippocampus but has minimal effects on axonal diameter and laminar counts

We obtained electron micrographs of the hippocampus, amygdala, prefrontal cortex (PFC) and nucleus accumbens and determined myelin sheath thickness, axonal diameter, and laminar count in ImageJ. In our primary analyses (males and females combined), the microbiome inoculation group had a significant impact (p-value <0.0125) on myelin sheath thickness in the hippocampus (F_3,11_ = 20.352, p = 0.0000853, three-way ANOVA). Post-hoc pairwise comparisons indicated that SPF mice had thicker myelin sheaths than HUM1, HUM2 and GF mice (Figure 6a). In addition, HUM2 mice had thicker myelin sheaths than HUM1 mice, though this result may reflect a single, influential individual. Sex-stratified analyses also suggested that microbiome composition may influence axonal diameter in the nucleus accumbens of male mice (F_3,10_ = 8.636, p = 0.00397, Two-way ANOVA). Post hoc tests did not indicate any significant pairwise differences, but visual inspection of results suggests HUM2 and SPF mice had greater axonal diameters in nucleus accumbens than HUM1 mice (Figure 6b). No other three-way ANOVA showed a significant effect for microbiome inoculation group (See Suppl data S5(a and b)). While our primary analyses did not indicate a significant interaction between microbiome inoculation group and sex for myelin sheath thickness, axonal diameter, or laminar count, sex-stratified analyses suggest the impact of gut microbiome inoculation group on myelin sheath thickness in the hippocampus may be stronger in males (F_3,8_ = 22.07, p = 0.000318, Two-way ANOVA) than females (F_3,3_ = 3.378, p = 0.172, Two-way ANOVA). Similarly, sex-stratified analyses also suggested that microbiome composition may influence laminar counts in the hippocampus (F_2,5_ =7.603, p = 0.030462, Two-way ANOVA) and amygdala of female mice (F2,5 =6.305, p = 0.043, Two-way ANOVA). In females, HUM2 mice had lower laminar counts than HUM1 mice in the hippocampus and lower laminar counts than GF mice in the amygdala (Supp S5 (c)). We also examined the myelin g-ratio. Three-way ANOVA did not reveal a main effect of microbiome inoculation group in any region but did indicate a significant interaction between sex and microbiome inoculation group for g-ratios in PFC. However, sex stratified analyses did not reveal significant effects of microbiome inoculation group in either males or females (Suppl S5(d)).

**Figure 6:**
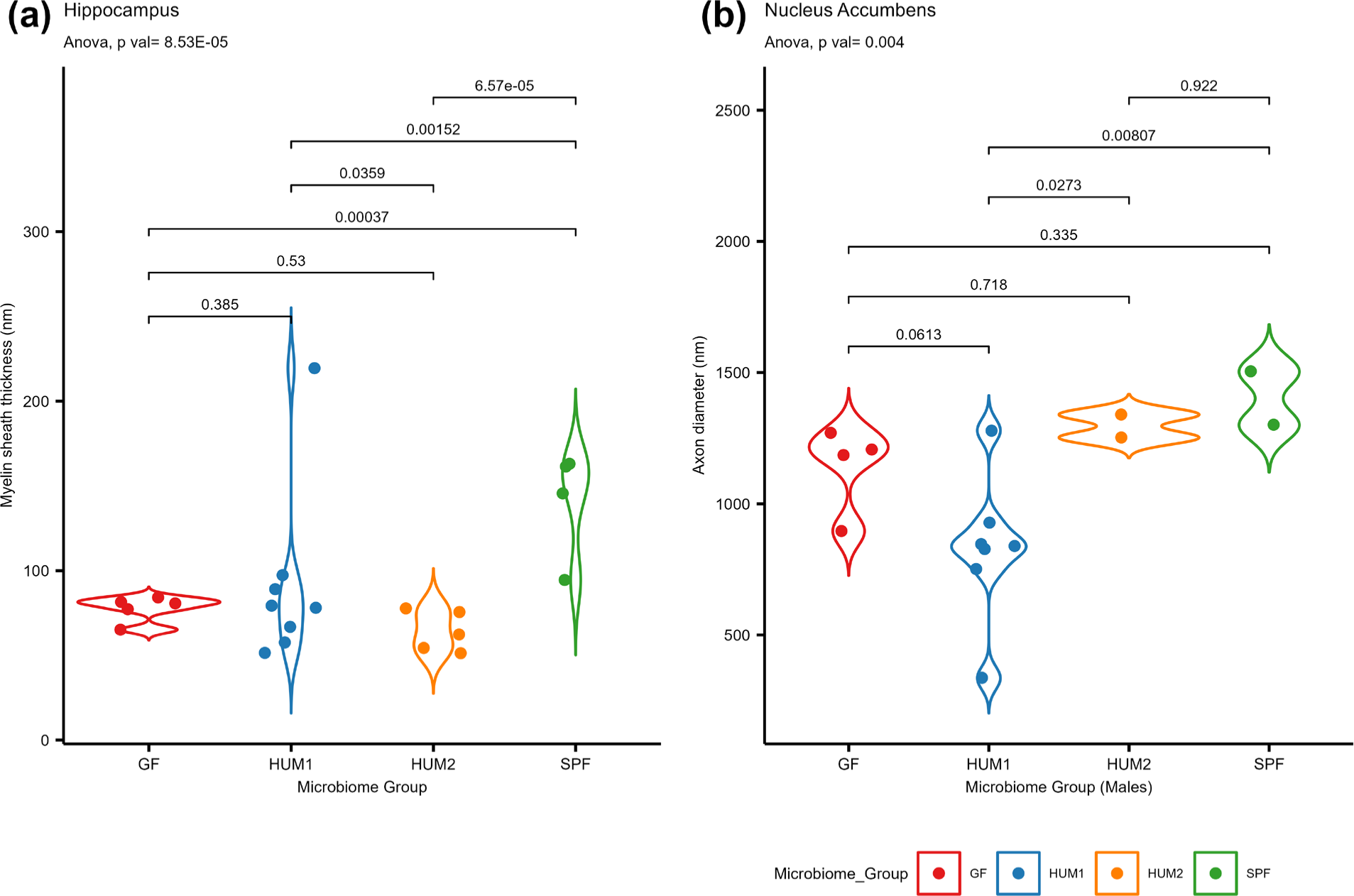
Effect of gut microbiome on myelination and axonal diameter, (a) myelin sheath thickness (nm) in hippocampus (cohort wise-M+F), (b) axon diameter (nm) in nucleus accumbens (male).

**Figure 6:**
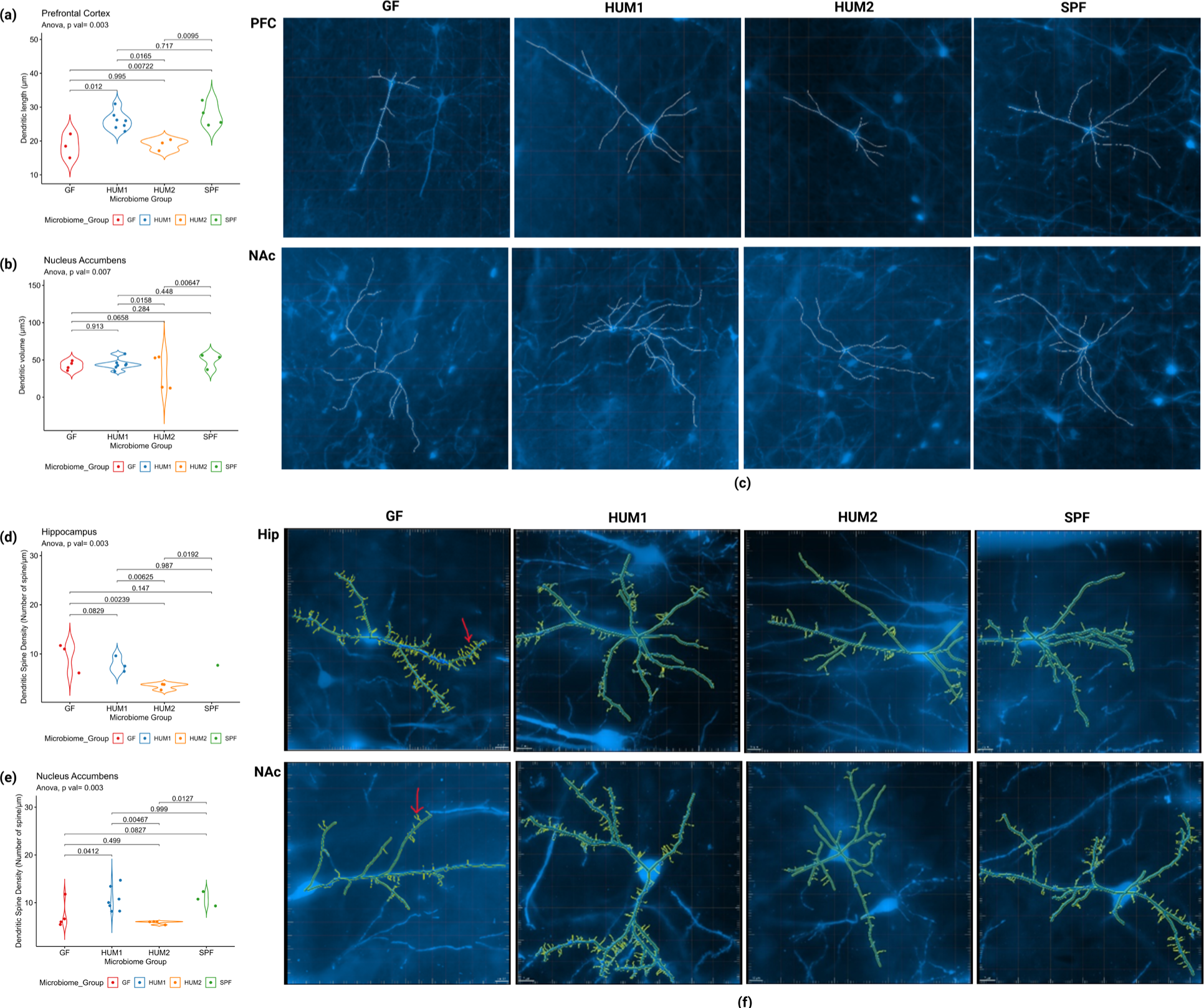
Effect of gut microbiome on dendritic morphology. (a) SPF and HUM1 had higher dendritic length in PFC, (b) SPF and HUM1 had higher dendritic volume in the Nucleus accumbens (NAc), (c) Representative photomicrograph dendritic length of GolgiCox stained neurons in the PFC and NAc, scale bar 20µm and examined with microscopy under 10X magnification. Spine density (d-f), (d) GF, HUM1 and SPF mice had more spine density in hippocampus (HIP), (e) HUM1 and SPF mice had more spine density in NAc, (f) Representative photomicrograph spine density of GolgiCox stained neurons in the HIP and NAc, scale bar GF and HUM2 10µm, HUM1 and SPF 7µm, and examined with microscopy under 40X magnification. Red color arrows point to examples of dendritic spines.

### Dendritic morphology differs between gut microbiome inoculation groups

Dendritic length, volume, and spine density were assessed in PFC, nucleus accumbens, amygdala and hippocampus using Golgi-Cox staining. Gut microbiome inoculation groups differed significantly (p-value <0.007) in dendritic length in PFC (F_3,7_ = 13.934, p = 0.00245, three-way ANOVA). Post-hoc analyses revealed that HUM1 and SPF animals had longer dendrites than GF and HUM2 animals (Fig 7a and 7c). Sex-stratified analyses suggest this pattern is present in both males (F_3,4_ = 7.09, p = 0.044, Two-way ANOVA) and females (F_3,3_ = 22.76, p = 0.0145, Two-way ANOVA). Primary analyses (combined sex) did not show significant effects of the microbiome inoculation group on dendritic length in the amygdala, nucleus accumbens, or hippocampus (supplementary data S6 (a)). No significant interactions with sex were identified. Sex-stratified analyses were conducted, but small sample sizes limit the conclusions that can be drawn (see supplementary data S6 (a)).

**Figure 7:**
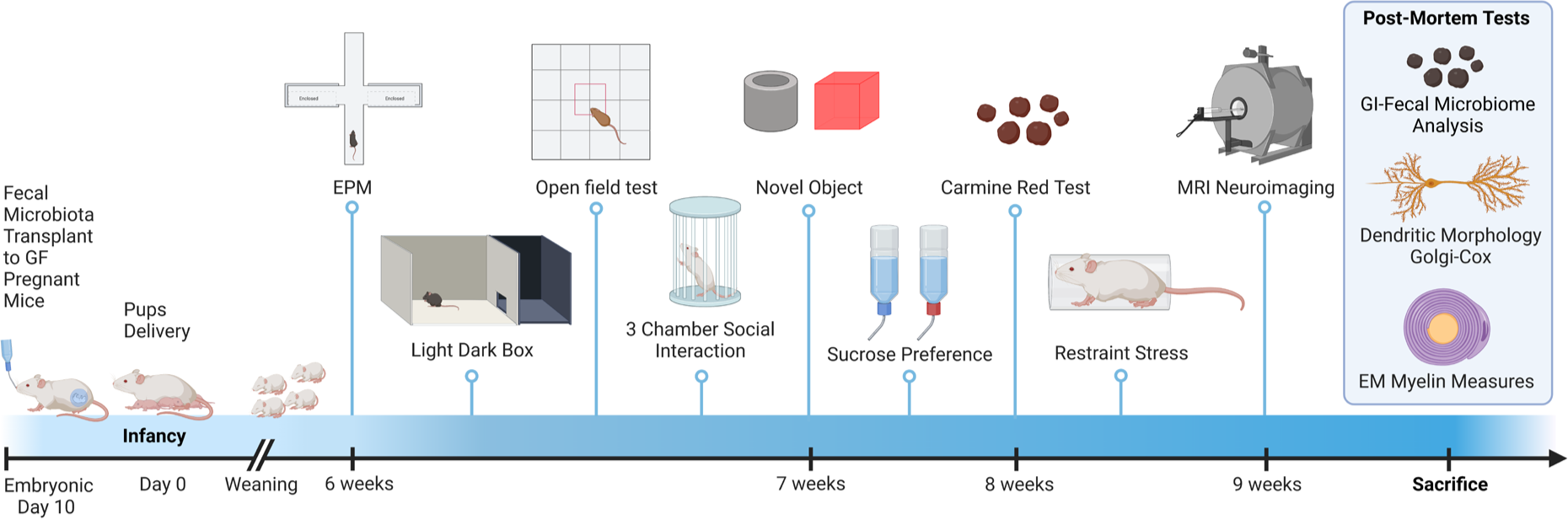
Schematic presentation of experimental procedure Donor phenotype/characteristics.

Microbiome inoculation groups differed significantly in the nucleus accumbens (F_3,8_ = 8.594, p = 0.006965, Three-way ANOVA) for dendritic volume, but not in the other three regions. HUM1 and SPF mice had more dendritic volume than HUM2 mice (Fig 7b and 7c). While our primary analyses did not indicate significant interactions between microbiome inoculation group and sex, exploratory sex-stratified analyses suggest potential sex-specific effects. Female HUM1 and SPF mice had more dendritic volume than female GF and HUM2 mice in nucleus accumbens (F_3,3_ = 114.7, p = 0.00136, Two-way ANOVA) (graphs included in supplementary data S6(b)). In hippocampus, male GF mice had higher dendritic volume than HUM2 male mice (F_2,2_ = 25.37, p = 0.0379, Two-way ANOVA, graph included in supplementary data S6(b)).

Combined sex analysis of spine density revealed an impact of microbiome inoculation group in hippocampus (F_3,3_ = 70.86, p = 0.00278, Three-way ANOVA) and nucleus accumbens (F_3,8_ = 11.008, p = 0.00327, Three-way ANOVA), but not in amygdala or PFC (see supplementary data S6 (c)). Post-hoc analyses indicate that in hippocampus, HUM2 mice have reduced spine density compared to all other groups (Fig 6d and 6f) while in the nucleus accumbens HUM2 mice have reduced density compared to both HUM1 and SPF mice and HUM1 mice have greater density than GF mice (Fig 6e and 6f), (F3,8 = 11.008, p = 0.00327, Three-way ANOVA). Sex-stratified analyses showed that female HUM2 and GF mice had lower spine density than HUM1 and SPF female mice in nucleus accumbens (F_3,3_ = 70.23, p = 0.00281 Two-way ANOVA, graph included in supplementary data S6(c)), while male mice did not show any differences in spine density (supplementary data S6(c)). However, in hippocampus male GF mice had higher spine density than HUM2 male mice (F_1,2_ = 403.3, p = 0.00247 Two-way ANOVA, graph included in supplementary data), while microbiome inoculation group did not have a significant effect on female mice (supplementary data S6(c)).

## Discussion

Studies of germ-free and antibiotic-depleted rodents provide abundant evidence that gut microbes influence brain development and the emergence of behaviors relevant to neuropsychiatric and neurodevelopmental disorders (*4*, *39*). There is also a growing literature demonstrating associations between early colonization patterns in human infants and behavioral outcomes including cognition (*23*, *40*), communication and social skills (*41*, *42*), temperament (*24*, *43–49*), hormonal responses to stress (*25*), regional brain volumes (*23*, *24*), and functional brain connectivity (*45*, *50*). However, these associations are not necessarily causal. We designed the present study to determine whether two microbial stool communities, commonly observed in human infants, impact behavior, myelination, dendritic morphology, and spine density, when introduced into mouse models. Like prior studies, we found reduced anxiety-related behaviors, increased locomotor/exploratory behaviors, and stronger corticosterone responses to an aversive situation in mice raised in germ-free environments as compared to SPF animals. Humanized animals were more like SPF mice than GF mice for most phenotypes, though both humanized groups exhibited reduced sociability in males. In addition, both humanized groups had thinner myelin sheaths in the hippocampus, as did GF animals. Humanized animals were similar to each other with the notable exception of dendritic morphology and spine density where mice in the HUM1 group had greater dendritic length in PFC, greater dendritic volume in nucleus accumbens, and greater spine density in both regions, compared to HUM2 animals.

Much of what is known about the influence of the gut microbiome on neurodevelopment comes from studies of mice raised without any exposure to microorganisms. The behavioral and physiological profile of these GF animals is well characterized and was reviewed in (*51*). GF mice tend to show decreased anxiety-like behavior (*22*, *52–55*), impaired memory (*52*), increased locomotor behavior (*22*), and a loss of the typical murine preference for social novelty (*56*), as well as increased corticosterone responses to stressors (*53*, *57*, *58*). Effects on social preference are mixed with some studies reporting increased social interactions with a conspecific (*55*), and others reporting social avoidance (*56*, *59*, *60*). Effects on depression-related behaviors, such as sucrose preference, a measure of anhedonia (i.e inability to experience pleasure) are also mixed with some studies reporting decreased depressive-like behavior (*61*– *63*), and others reporting no effect (*64*). Most of our results are in keeping with this prior literature. However, we did not observe altered social behavior or impaired memory in our GF mice, and our findings for depressive-like behavior suggest greater anhedonia in GF mice rather than less. One potential explanation for these discrepancies is that behavioral testing took place after our GF animals were removed from the GF facility in which they were raised. Assessments of their cecal contents at sacrifice suggest rapid colonization by members of class Clostridia and order Erysipelotrichales. Some aspects of the behavioral profile in GF mice are known to be reversible with bacterial colonization, including the lack of preference for novel versus familiar conspecifics (*56*), and this could explain our results.

Regarding behavioral alterations in humanized mice, our study is one of the first to evaluate the effects of different gut microbiomes, collected from human infants, on long-term behavioral outcomes in mice. Other studies have compared the effect of FMT from preterm and term neonates (*65*), preterm infants with and without normal weight gain (*66*), antibiotic-exposed and non-exposed infants (*67*), infants born to mothers with or without inflammatory bowel disease (*68*), infants with and without cow’s milk allergy (*69*), and healthy vs severely stunted and underweight infants (*70*). However, these studies focused on growth, inflammatory activation, and intestinal and metabolic abnormalities rather than behavior. Our study is also unique in looking at natural variation between typically developing infants who were exclusively breastfed, vaginally delivered, and unexposed to antibiotics, though we note a prior study did document that transplanted healthy infant microbiota mainly composed of *Bifidobacterium* and *Bacteroides* had a protective impact on sensitization and food allergy in mice (*71*). While our humanized animals were usually more similar to the SPF mice than to GF mice, we observed reduced sociability in humanized males regardless of whether their dams had been inoculated with human infant communities rich in *Bifidobacterium* or those rich in *Bacteroides*. This suggests neither community is sufficient to promote development of typical social behavior in male mice. Indeed, given that male mice raised in a GF environment were similar to SPF mice, it’s possible that these particular human infant communities actively disrupt the development of typical social behavior in male mice. The lack of a differential effect of the two human infant microbiomes on behavior contrasts with earlier studies comparing human microbiomes from children with and without ASD (*72*, *73*), and individuals with and without depression. For example, Sharon et al. (2019) demonstrated that transplanting microbiomes from ASD patients with ADOS scores between 6-10 to mice induced ASD-like behaviors in offspring that were not present in offspring colonized by microbiomes from children without ASD or those with mild ASD (ADOS scores from 4-5). This effect was more pronounced in male offspring and included decreased locomotion and decreased communication, but no differences in the three-chamber sociability test (*72*). Xiao et al (2021) reported increased anxiety, decreased sociability, and decreased preference for social novelty in GF mice receiving microbiomes from ASD children around 4 weeks of age as compared to mice receiving microbiomes for children without ASD. In a similar manner, transplant of microbiota from patients with depression induces anxiety and depression-related behaviors in mice that are not present in mice receiving microbiota from healthy, non-depressed individuals (*63*, *74*). Differences between the current study and these prior studies likely reflects differences in the age at which samples were collected from the human subjects. In addition, microbiomes from subjects with a clinical diagnosis of ASD or depression may be more frankly dysbiotic than samples from different, typically developing infants. Thus, future behavioral studies in this area ought to examine a wider range of infant microbiomes including ones disrupted by factors such as c-section delivery and antibiotic exposures. A broader array of outcomes could also be examined, including impaired communication and repetitive behavior, which are very relevant to ASD.

We also analyzed the effect of different gut microbiomes on axon diameter and myelination in the hippocampus, amygdala, PFC, and nucleus accumbens. Evidence for a link between the microbiome and developmental myelination was recently reviewed in *Neuroscience Letters* (*75*), and began with a seminal study by Hoban et al. (*6*). Hoban et al (2016) reported increased transcription of myelin regulatory and myelin component genes in PFC of GF mice at 10 weeks of age resulting in higher myelin sheath thickness in PFC axons (*6*). They did not observe similar changes in cerebellum, amygdala, hippocampus, striatum, or frontal cortex. An independent group also reported increased transcription of myelin-related genes in the adult PFC (6-8 weeks of age) following antibiotic treatment in infancy (*76*). However, Lu et al (2018, (77)) reported somewhat opposing findings with GF mice demonstrating reduced myelin content in corpus callosum, anterior commissure, fimbria, and internal capsule at 4 weeks of age and no differences in frontal or parietal cortex, hippocampus, striatum, or thalamus. The only difference that persisted until 12 weeks of age was reduced myelin content in the internal capsule. We found that GF mice had lower myelin sheath thickness in hippocampus compared to SPF mice, findings which are more like those of Luo et al (2018, (*77*)) than those of Hoban et al. (*6*) and Keogh et al. (*78*). This could be due to rapid colonization of our GF animals by members of class Clostridia and order Erysipelotrichales, as Hoban et al. found that colonization with a conventional microbiota following weaning reversed changes in myelin and activity-related gene expression. It could also reflect methodological differences between the various studies. We also observed reduced myelin sheath thickness in the hippocampus of HUM1 and HUM2 mice. Interestingly, transplant of microbiota from preterm infants with poor growth reduces cortical MBP expression at 2 weeks of age when compared to transplant from high-growth infants (*79*), suggesting certain kinds of infant gut microbiomes disrupt oligodendrocyte development in mice, and this could explain our results. However, we cannot rule out the possibility that microbiome composition later in life was more critical to our findings. Future studies could sacrifice animals at different ages to resolve this question. Future studies could also investigate potential mechanisms by which our human microbial communities altered myelin sheath thickness. The study of preterm infants suggests neuroinflammation and circulating IGF-1 may play a role, while Lu et al (2018, (*77*)) highlight butyrate as a potential mechanism. Regardless of mechanism, reduced myelin sheath thickness in the ventral hippocampus is expected to reduce the speed of electrical transmission and could influence behaviors linked to this region (*80*). Though we did not observe altered anxiety-related behaviors in our humanized rodents, reduced myelin sheath thickness might contribute to the altered social behavior seen in our humanized male rodents, especially since a circuit connecting the basolateral amygdala (BLA) and ventral hippocampus (vHPC) is involved in the modulation of social behavior in the three-chamber test (*81*).

Perhaps the most intriguing findings of the current study were the ones for dendritic morphology. We found that the HUM1 group had greater dendritic length in PFC, greater dendritic volume in nucleus accumbens, and greater spine density in both regions, compared to HUM2 animals, suggesting this particular community may promote greater local connectivity. The PFC findings are especially intriguing as our group previously reported that human infants with higher abundance of *Bifidobacterium* at one month of age have higher volumes of PFC at one year of age (*24*), which could reflect greater dendritic complexity. We did not include the nucleus accumbens in that prior study, but given the current results, it will be included in our future human work. Prior studies have reported GF mice exhibit dendritic hypertrophy of BLA aspiny interneurons, increased dendritic length and spine density of BLA pyramidal neurons, dendritic atrophy and reduced spine density on ventral hippocampal pyramidal neurons (*82*), and dendritic elongation of pyramidal neurons in the anterior cingulate (*83*). We did not replicate these findings in our GF mice. This could be due to rapid colonization of our GF animals by members of class Clostridia and order *Erysipelotrichales*, as discussed previously. Nevertheless, our results add to a growing body of evidence that variation in the gut microbiome influences dendritic phenotypes. The key question that remains is why the two infant microbiomes have differential effects on dendritic development. 16S rRNA gene sequencing of post-mortem colonic contents indicated that membership in the HUM1 group was predicted by higher relative abundance of *Turicibacter, Bifidobacterium, Candidatus Arthromitus,* and *Erysipelotrichaceae*, while membership in HUM2 was predicted by higher relative abundance of *Bacteroides, Lachnospiraceae_NK4A136_group, Anaeroplasma, Clostridia_vadinBB60_group, Parabacteroides, Papillibacter* and *Flavonifractor.* These genera may have differential metabolomic and/or inflammatory profiles that either promote or inhibit dendritic development. Notably, a recent study showed that oral administration of probiotics containing *Bifidobacterium* increases spine density in hippocampus (*84*). Another study revealed that administration of *Bifidobacterium* enhances spine density in pyramidal neurons in rat hippocampus (*85*). Furthermore, infant-associated *Bifidobacterium* has immune modulatory and protective properties critical for immune education, while *Bacteroides* does not (*86–88*). These findings suggest higher abundance of *Bifidobacterium* may contribute to dendritic morphological changes in HUM1 animals. However, we also want to note that *Turicibacter* can induce serotonin production in the gut, which may indirectly influence neurodevelopmental process (*89*). Another question that remains is whether these structural changes have functional consequences not captured by our behavioral tests. Dendritic arbors receive most synaptic inputs while dendritic spines represent the most important parts of excitatory synapses in the hippocampus, neocortex, and other brain regions (*90–92*). Consequently, the alterations we observed could affect synaptic transmission and plasticity such that HUM1 mice have a more active PFC and nucleus accumbens than HUM2 animals. Our PFC slices came from anterior cingulate cortex, as defined by Heukelum et al (*93*), which also includes prelimbic areas. This region regulates autonomic functions, depressive-like behaviors, reward, negative affect, and pain (*93*). The nucleus accumbens is part of the mesolimbic pathway. It is critical for motivated behaviors and reward processing and is thought to play a role in the pathogenesis of major depression (*94*) and addiction (*95*). While we did not observe differences between HUM1 and HUM2 animals in the post-hoc tests for sucrose preference, there was a main effect of microbiome inoculation group on sucrose preference with HUM1 animals being more similar to SPF animals and HUM2 showing lower preference for sucrose, albeit not as low as the GF animals. Given the results of our dendritic complexity analyses, it would make sense to include more behavioral measures relevant to depression in future studies, such as the forced swim test, and to investigate addictive behaviors. In the future, we also plan to analyze the impact of gut microbiome on different types of spines including filopodia, mushroom, thin and stubby spines.

The current study has both strengths and weaknesses. Strengths of the study include the comprehensive behavioral assessments, use of electron microscopy to evaluate axonal phenotypes, and use of Golgi-Cox to achieve high-quality structural stains of neurons. We have also endeavored to follow Guidelines for Reporting on Animal Fecal Transplantation (GRAFT; (*96*)), to improve rigor and reproducibility (See Supplementary Table S1). We also note that the mouse model is preferred over pig models for transplantation studies of human infant microbiomes and that *Bacteroides* colonizes particularly well in this system (*26*). That being said, human microbiomes undergo substantial changes when introduced into mice, in part because some microbes have host-adapted features that limit their ability to colonize other species (*26*, *97*, *98*). This is a general limitation of humanized mouse studies. Some microbes do not colonize at all and others, including *Bifidobacterium*, colonize at much lower abundances compared to human donors (*26*, *97*, *99*). We observed this in the current study where the abundance of *Bifidobacterium* in the inocula was around 70%, but the abundance in post-mortem colonic contents in HUM1 animals was around 1 to 4%. Future studies could attempt to increase the engraftment of *Bifidobacterium* by maintaining anerobic condition during inocula creation, performing multiple inoculations, and/or by supplementing the mouse diet (*100*, *101*) to promote colonization and survival of this group. We also acknowledge general limitations of using GF mice. GF mice are often preferred for microbial transplantation studies, as competition between resident and newly transplanted microbes is absent. However, the absence of microbiota in GF rodents produces immune system impairments (*102–105*), metabolic alterations (*22*, *102*, *106*), and differences in social behavior (*56*). Importantly, even after recolonization of the gut microbiome these alterations are not completely reversed (*56*, *103–105*). Our results might have been different if we had transplanted our microbiomes into antibiotic depleted dams rather than GF dams. Another limitation was the use of 16S rRNA gene sequencing rather than metagenomic sequencing to assess donor communities, inocula, and engraftment in the mouse models. While 16S rRNA sequencing is a well-established approach and has been the major driver of microbiome research over the last several decades, shotgun metagenomic sequencing has several advantages. This includes 1) direct inference of relative abundance of microbial functional genes; 2) species- and strain-level resolution for known organisms; 3) capturing phages, viruses, and microbial eukaryotes in addition to bacteria, and 4) identification of novel gene families (*107*). Furthermore, 16S rRNA gene sequencing may underrepresent some phyla as compared to metagenomic sequencing (*108*, *109*), perhaps due to unequal amplification of different 16S rRNA genes (*110*). We also note that recent research suggests the fecal microbiome may be regulated by circadian rhythms in the host (*111*), but we did not record the time of collection for human fecal samples. We will consider all these limitations in future studies.

In conclusion, the current study provides novel information about the impact of different gut microbiomes, including those from human infants, on behavior, myelination, and dendritic morphology in mouse models. Results add to a growing body of literature suggesting that the gut microbiome influences brain development, which could contribute to risk for neurodevelopmental and psychiatric disorders. While humanized mouse models have limitations, they also hold great potential for testing early interventions that could promote optimal development of the gut microbiome and improve behavioral outcomes.

## Materials and Methods

### Overview

The objective of the current study was to determine if variation in human infant gut microbiomes influences neurodevelopment using humanized mouse models. We transplanted microbial communities derived from human infants into pregnant germ-free mice and evaluated offspring behavior with well-established paradigms for anxiety-related, depression-related, exploratory, and social behaviors, as well as object memory. Cortisol reactivity to restraint stress and intestinal transit time (Carmine red test) were also measured. After animals were humanely euthanized, we analyzed cecal contents using 16S rRNA sequencing, assessed dendritic morphology and spine density using Golgi-Cox staining, and assessed axonal and myelin-related phenotypes using transmission electron microscopy. A subset of animals (see Suppl data S7) underwent MRI; those results are still being processed and will be the focus of a separate manuscript. The experimental procedure is schematically shown in Figure 7. Fecal transplantation and subsequent tests were conducted in 3 cohorts across a period of approximately 11 months (August 2020 through June 2021).

83 human infant samples were collected as part of an NIH-funded study on gut microbiome and anxiety-related behaviors. The institutional review board (IRB) of the University of North Carolina at Chapel Hill approved the research protocol (protocol #14-0932, MSU reliance agreement STUDY00001259), and written informed consent was obtained from a parent/legal guardian of each subject. Inclusion criteria for participation in the study were vaginal delivery and exclusive breastfeeding until the first study visit at 1 month. Participants were excluded for maternal antibiotic usage two weeks before delivery (including Group B Streptococcal prophylaxis), antibiotics given to the infant before the first study visit, neonatal intensive care unit stay, birth weight <2500 g, gestational age <37 weeks, major maternal medical illness, prenatal drug use, primary language other than English, and fetal ultrasound abnormalities. To identify groups of subjects with similar microbial communities, we conducted 16S rRNA sequencing of the V1-V2 gene region followed by amplicon sequencing data analysis with the Quantitative Insights Into Microbial Ecology (QIIME) software pipeline (*112*). Unweighted UniFrac distance, weighted UniFrac distance, Bray-Curtis distance (BC), Jensen-Shannon Divergence (JSD), and the square root of JSD (rJSD) were applied to the relative genus abundance in each individual. The Calinski-Harabasz (CH) Index, silhouette index (SI), and prediction strength (PS) were used to determine the optimum number of clusters (Figure 8, panels a-d). Two clusters were identified. One cluster was characterized by relatively high levels of *Bacteroides* and the other by relatively low levels of this genus (Figure 8e). Samples in the latter cluster were often characterized by a high abundance of a particular taxa, but the identity of that taxa varied across infants. Some had very high abundance of *Bifidobacterium*, some had very high abundance of *Streptococcus*, and some had very high levels of *Clostridiales*. For the current study we focused on infants with relatively high abundance of *Bifidobacterium* (HUM1) and those with a relatively high abundance of *Bacteroides* (HUM2). 3 donors per group (HUM1, HUM2) were selected from the sample of 83 infants (Figure 8e). All donors were white and non-Hispanic. HUM1 donors included 1 male and 2 females; all HUM2 donors were male. HUM1 donors were between 40 and 72 days of age; HUM2 donors were between 40 and 51 days of age.

**Figure 8.**
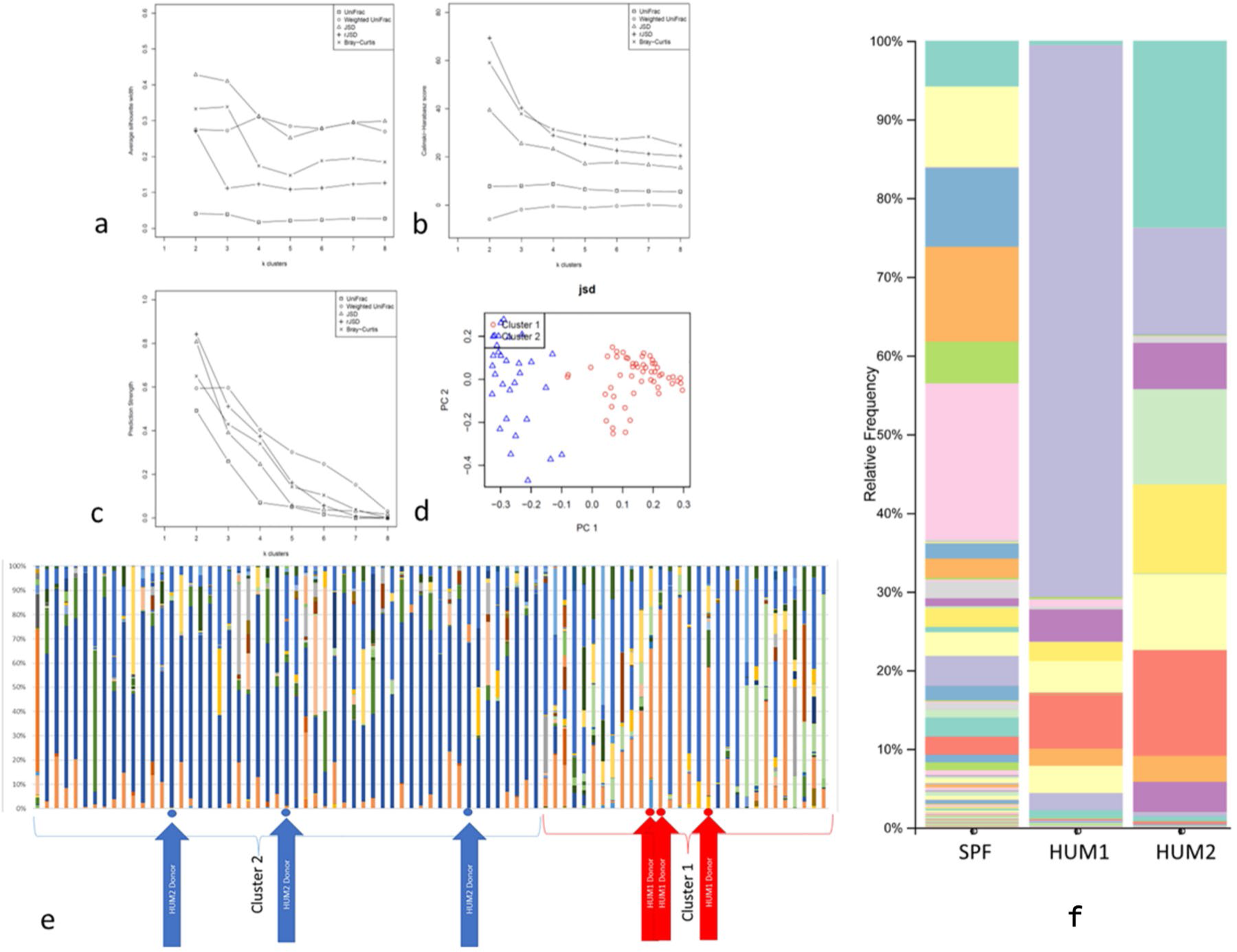
Selection of human donors. **(a-e)**. Distance metrics including unweighted Unifrac, weighted Unifrac, Jensen-Shannon, Root Jensen-Shannon, Bray-Curtis were scored with average silhouette width (**a**), Calinski-Harabasz (**b**), and prediction strength (**c**) to identify the optimal number of clusters. Most measures supported a two-cluster solution. (**d**) Principle coordinate plot with sample color coded by cluster (Cluster 1 in red, Cluster 2 in blue. (**e**) Infant samples sorted by cluster. Cluster 2 samples are typically dominated by *Bacteroides* (dark blue), while Cluster 1 samples have low abundance of that genus. Samples in Cluster 1 are often dominated by *Bifidobacterium* (medium orange) *Streptococcus* (light green), or *Clostridium* (medium blue). Donor samples are indicated by red (HUM1) and blue (HUM2) arrows. (**f**) **Taxonomic composition of fecal inoculums.** The most abundant taxon in the HUM1 inoculum is *Bifidobacterium* (lilac). The most abundant taxon in the HUM2 inoculum is *Bacteroides* (aqua). The most abundant taxon in the SPF community is an unnamed genus in the family *Lachnospiraceae* (pink).

To create the mouse SPF and autoclaved SPF inoculums, we collected fecal pellets from Swiss Webster SPF mice between 6-8 weeks of age. 8 mice (4 males and 4 females) were used. All SPF donor mice were co-housed, and cages were kept on adjacent racks. Animals were fed with autoclaved water in glass water bottles and autoclaved food (LabDiet 5013), under standard housing condition (temperatures at 72 ± 2 °C, 30-70% humidity and light cycle 12h:12h). The animals in our study were shown to be free of pathogens including mouse hepatitis virus, minute virus of mice, mouse parvoviruses, murine encephalomyelitis virus, mouse rotavirus, Sendai virus, pneumonia virus of mice, mouse reovirus, mouse adenoviruses, lymphocytic choriomeningitis virus, ectromelia virus, polyomavirus, mouse cytomegalovirus, mouse thymic virus, CAR bacillus, Mycoplasma pulmonis, Encephalitozoon cuniculi, Corynebacterium bovis, murine ectoparasites (Myocoptes, Myobia, Radfordia) and mouse pinworms (Aspiculuris tetraptera, Syphacia obvelata).

#### Donor Sample Collection and Storage

Participating families were mailed a sample collection kit shortly before an in-person study visit. Parents were instructed to collect 3 scoops of fecal material (∼665 mg of feces) using the spoon attached to the cap of a 16.5ml specimen tube (Sarstedt Inc, Catalog # NC0729971). Samples were collected from a single soiled diaper and immediately placed in a tube completely submerged in 10 ml of thioglycolate broth (BD Difco™ Fluid Thioglycollate Medium Catalog # 225650), a multipurpose, enrichment medium that produces a range of oxygen concentrations along its depth. This makes it suitable for supporting the viability of obligate aerobes, obligate anaerobes, facultative anaerobes, microaerophiles, and aerotolerant organisms. Before sending the thioglycolate broth to parents for fecal sample collection, it was sterilized by autoclaving for 20 min at 121 °C and aliquoted into sterile tubes. Parents placed the sample tube into a biohazard bag and recorded the date of collection. Time of collection was not recorded. They then placed the bag into a Styrofoam container atop a freezer pack and placed the container into their home freezer. Parents were instructed to wear non-latex hospital gloves (provided in their study kit) to minimize contamination. Samples were brought to the UNC lab within 24 hours of collection. Once received, the tubes were stored at −80 °C until they were shipped to Michigan State in June of 2020. Samples were shipped overnight on dry ice and all samples were confirmed to be still frozen on arrival. They returned to −80 °C storage until the inoculums were prepared on 08/27/2020. In total, time from collection to creation of the inoculums ranged from 26-39 months. To protect the samples from degradation, we avoided multiple freeze-thaw cycles. Parents also provided ∼200 mg of feces in a separate tube containing 1 ml Allprotect reagent (Valencia, CA), which was used for 16S rRNA sequencing.

Mouse fecal samples were collected between 9:00 am to 12:00 pm. Mice were taken from their cage by holding the tail base and placed over the food rack of a clean cage. After that the researcher slightly lifted the back portion of the mouse body by holding the base of the tail and put marginal pressure by finger over back near to tail and collected the freshly defecated fecal pellets (1–2) in a sterile 1.5 ml Eppendorf tube. To avoid contamination, the fecal sample was collected under a biosafety hood and the collector wore PPE including a mask and nitrile gloves. Immediately after collecting the fecal pellets, inoculums were prepared at room temperature as described below.

#### Creation and storage of inoculums

All human fecal samples were processed at room temperature under a biosafety cabinet. An equal volume of 20% glycerol in phosphate buffer solution (PBS) was added to the original stock then mixed using a pipette for 2 minutes. The resulting fecal slurry was divided into 2 ml aliquots. The final concentration of human fecal inoculum was ∼227mg/ml. For the SPF inoculum, ten fresh fecal pellets (400mg) were collected from 4 male and 4 female mice (1-2 pellets from each mouse) then added to 10 ml PBS plus 10% glycerol solution under room temperature conditions in a Biosafety cabinet. A TissueRuptor (Qiagen) was used to homogenize the sample at 30000rpm for 60 sec at room temperature before dividing into 1 ml aliquots, each in a cryotube (ThermoFischer). The final concentration of mice fecal inoculum was 40mg/ml. Half of the aliquots were autoclaved before freezing.

16S rRNA amplicon sequencing was used to assess quality and composition of the inocula (1 aliquot per group). Full details on sequencing methods are provided in a later section. The outcome of the quality assessment was relative abundance of different taxa as shown in Figure 8(f). Similar to the original donor samples, the HUM1 inoculum had a very high relative abundance of *Bifidobacterium* (70.12%), which is similar to the donor samples. The most abundant taxa in the HUM2 inoculum is *Bacteroides* (around 23.75%). This is lower than the donor samples which had *Bacteroides* abundance ranging from around 60% to around 80%, suggesting that this group was negatively impacted by collection, storage, and/or processing procedures. The most abundant taxon in the SPF community is an unnamed genus in the family *Lachnospiraceae* (19.58%), (Suppl S-1(j and k)). Inoculums were stored at −80 °C at MSU, then transported on dry ice to the University of Michigan Germ-Free Mouse Facility on the day of administration and returned to −80 °C conditions. Samples were kept in the port of the isolator and the tubes sprayed with sterilant before administration to the mice. After one-hour, thawed samples in the isolator were administered to the pregnant female mice in that isolator. For practical reasons, anerobic conditions were not used during fecal sample collection, transport, processing, and storage neither for mice fecal nor for human samples. The total time from creation of the inoculums to administration to mice ranged from 1-6 months.

#### Recipient Animals and Fecal Transplant Procedure

Pregnant Swiss-Webster germ-free (GF) mice were generated by the University of Michigan Germ-Free (GF) Mouse Facility. We chose Swiss Webster mice because they have been widely used in murine microbiome studies, facilitating comparisons to the existing literature, have relatively large litters, lactate well, and show strong mothering abilities. All recipient mice (2-3 mice/group) were between 6 and 8 weeks of age at the start of pregnancy and were randomly selected for fecal slurry inoculations. The GF Mouse Facility stores animals in flexible film soft-sided germ-free isolators and monitors for potential contamination events by routine sample testing. Recipient animals were group housed prior to mating. After mating, staff checked all mated females for vaginal plugs, separated all males, and regrouped females into social housing (2-4 females/cage). Females were moved to single-housing on day 10 of pregnancy and cages were arranged next to each other with a specialized positive-pressure individually ventilated rack system. No cage was exposed to anything from any other cage on the rack; cages are completely sealed and airflow in and out was controlled through several filters. Each cage acted as a miniature single isolator. Each cage used sterile bedding (Tek-fresh) and enrichment (Enviropak), autoclaved water in glass water bottles and autoclaved food (LabDiet 5013), under standard housing condition (temperatures at 72 ± 2 °C, 30-70% humidity and light cycle 12h:12h).

Fecal slurries were administered by oral gavage on embryonic day 10 (ED10) between 1:00-2:00pm as a single dose of 200µL. Group 1 (HUM1 group) was inoculated with a mixed human fecal slurry derived from infants with relatively high levels of *Bifidobacterium*. Group 2 (HUM2 group) was inoculated with a mixed human fecal slurry derived from infants with relatively high levels of *Bacteroides*. Group 3 (SPF control) received fecal slurry from specific pathogen free mice (SPF group) and Group 4 (GF group) received autoclaved fecal slurry from SPF mice. Recipient pregnant mice were not fasted, anesthetized, or manipulated in any other way prior to gavage. Gavage was done with manual restraint and metal gavage bulbs. There were no adverse events observed in recipient pregnant mice in response to fecal slurry inoculation. The University of Michigan GF facility monitors the germ-free condition of animals by performing periodic bacterial growth assays from fecal samples and moist chow (every 4-8 weeks) and 16S rRNA PCR fecal test twice a year. Cages are also monitored with a “mold trap” as a visual indicator for sterility. GF facility confirmed that all animals were in GF status before fecal slurry transplant and GF group animals remained in GF conditions until transferred to MSU. For the first three groups (HUM1, HUM2, and SPF) pregnant mice were transferred to the Michigan State University Animal Care Facility between Day 14 and Day 20 of pregnancy in sterile containers. The GF group remained at University of Michigan through weaning and the offspring were transferred to MSU around 5 weeks of age.

#### Housing Conditions at Michigan State University

Campus Animal Resources (CAR) provided veterinary care, daily husbandry, and health checks for all animals. Animals were housed in a special room dedicated to mice carrying human microbiotas. To prevent cross-contamination between groups of mice with different microbiotas, cage changes were carried out in a Biosafety cabinet. Gloves worn for cage changes were decontaminated with CAR supplied disinfectant solution (Virkon(TM)S tablet) between each cage and changed completely between each group of cages. Food (Teklad Global 2019S), water, cages, and bedding (Bed-o’Cobs 1/8”) were autoclaved. Optimice® IVC rack system and Optimice® mouse IVC cages were used. Cages were arranged in columns such that every column included only one group of animals and an empty column was kept between each group. Each cage had a separate ventilated rack system, no cage was exposed to anything from any other cage on the rack. Each litter occupied a single cage, with their mother, until weaning. After weaning animals were group housed with littermates of the same sex (4-5 animals per cage) until the sucrose preference test. Animals were singly housed throughout the sucrose preference test, carmine red test, and the stress reactivity test. They remained singly housed until the end of the study.

### Behavioral studies

Both male and female offspring were studied (Table 1) Table 1. Treatment groups of animals with sex details Offspring were assessed between 6-9 weeks of age with various tests including elevated plus maze (EPM) (day 1), light/dark preference (day 2), open field (day 3), and three-chamber sociability tests (day 4 or 5) in week 6. In week 7, we performed novel object recognition (day 1 or 2) and the sucrose preference test (day 3-5). In week 8, we performed carmine red testing (day 1 or 2) and the stress reactivity test (day 4 or 5).

**Table 1.** Treatment groups of animals with sex details.

| Group | Number of animals |  |  |
| --- | --- | --- | --- |
|  | Male | Female | Total |
| HUM1 | 13 | 6 | 19 |
| HUM2 | 4 | 7 | 11 |
| SPF | 4 | 4 | 8 |
| GF | 6 | 5 | 11 |

All behavioral testing except the sucrose preference and carmine red tests were conducted in a dedicated quiet behavior room and performed between 9:00am to 6:00pm. All animals were transferred to the behavior testing room 30-45 min prior to beginning each paradigm to habituate to the condition of the behavior testing room (*28*). After each trial, all the urine and fecal boli were removed. The tester disinfected the behavioral apparatus and their gloved hands with 70% v/v alcohol before and after handling each animal. GF group animals were handled first, followed by HUM1 group animals, then HUM2 group, then SPF. Only one microbiome inoculation group was allowed in the testing suite at any given time and PPE were changed completely between each group of mice. Animal movement was tracked and processed by TopScan camera and software (Clever Sys Inc., VA).

#### Elevated plus maze

The elevated plus maze is made of dark gray acrylic plastic, with two open and two closed arms, The open arms are across from each other and perpendicular to the two closed arms. The dimension of each arm was 28×5cm and the central area was 5×5cm. The entire apparatus was raised above the ground to a height of at 50cm. Two arms were open, and two arms were closed with 15cm high walls made of same material. Each mouse was placed in the center area of the maze with its head directed toward a closed arm and allowed to move freely about the maze for 10 min. The following behaviors were recorded and assessed: number of entries in open arm, distance traveled in open arm, time spent in open arm, number of entries in closed arm, time spent in closed arm, distance traveled in closed arm time spent in central area, distance traveled in central area, number of entries in central area, total number of entries in open arm and closed arm, and percentage of entries in open arm (*27*, *28*).

#### Light/dark preference test

The apparatus used for the light/dark transition consisted of a cage 40 (width) x 35 (height) x 40 (depth) cm divided by a 35 cm tall partition with a door. One chamber was brightly illuminated (200-400 lux), whereas the other chamber was dark (5 lux or less). The mouse was placed into the middle of the light chamber and allowed to move freely between the two chambers for 5 min. Time spent in light chamber and dark chamber, and number of entries into the dark chamber were recorded (*29*, *30*, *113*).

#### Open field test

Each animal was placed in the center of an open field box (27×27×27cm) with high walls to prevent the animal from escaping the apparatus. The apparatus floor has gridlines. These gridlines divided the area into equal smaller squares allowing software to determine when the mouse is in the outer versus the inner zone. Four mice were tested at the same time, each in a different apparatus (one mice per apparatus). Animals were allowed to explore freely for 5 minutes. Using the Topscan software, we calculated time spent and distance traveled in the inner and outer zone. In addition, the experimenter counted the number of fecal boli present at the conclusion of the trial (*32*, *114*).

#### Three-chamber sociability test

Sociability and preference for social novelty were analysed using a polycarbonate rectangular three chamber apparatus, having dimension 60 x 40 x 25 cm. Each side chamber connected with the middle chamber with a sliding door (4 x 4 cm), to allow free movement of the animal between the chambers. Testing was conducted in four phases. In phase 1, the test mouse was allowed to habituate to the central chamber for 10 minutes. In phase 2, the test mouse could explore all 3 chambers freely for 10 minutes. In phase 3 (sociability session), we introduced a stimulus (Stranger I) mouse (non-littermate control SPF mice with same gender but housed separately) in a small wire cage in one side chamber and an empty wire cage in the other side chamber and allowed the test mouse to explore freely for 10 minutes. Sociability was evaluated by measuring the time spent by test mice in the chamber with the Stranger I mouse and the time spent in the chamber with the empty wire cage. In phase 4 (social novelty session), we introduced a second stimulus mouse (Stranger II) into the empty wire cage and allowed the mouse to explore freely for 10 minutes. Outcomes for Phase 2 include time spent with the familiar mouse (Stranger I) and novel mouse (Stranger II) during the social novelty session and the discrimination Index (*77*).

#### Novel object recognition

Novel object recognition was performed as described by Lueptow (2017)(*115*), using the same apparatus/ arena used for the open field test (a square chamber 27×27×27cm). The test was conducted in three phases. In phase 1, mice were habituated to the empty arena for 5 min. In phase 2, the experimenter placed two identical objects (white cubes) in the NE and SW corners of the arena and returned the mice to arena to explore for 5 min. After completion of phase 2, animals were returned to their home cage for an hour at which point Phase 3 commenced. During phase 3, the mouse was allowed to explore the empty arena for 5 minutes. Then a familiar object (white cube) and a novel object (grey cylinder) were placed in the NE and SW corners. Mice continued to explore for 5 min. Exploration of an object was defined as nosing and sniffing at the object or being in very close proximity to the object (within 2 cm). Outcomes are based on the final phase of the experiment and total time spent exploring the familiar object, total time spent exploring the novel object, % recognition index [time spend to explore novel object/ (time spend to explore novel + familiar object) X 100] (*33*, *115*).

#### Sucrose preference test

The sucrose preference test was performed as described by Iñiguez et al., 2009 (*116*). This test is used to assess anhedonia in mice. Anhedonia is a common symptom of depression in humans. Animals were moved to single housing and each cage was fitted with two sipper bottles. One bottle was filled with water and other with 1% sucrose solution. Mice can choose to consume water or sucrose. Potential side preference bias for bottles were counterbalanced by switching the bottle position once a day. We monitored liquid intake by weighing each bottle daily for 4 days, then calculated % preference of sucrose over water [sucrose/(sucrose+water) X100] (*35*, *117*).

#### Carmine red test

Carmine red tests were performed as described by Dey et. al. (2015) to determine gut transit time. Carmine red (Sigma-Aldrich) was dissolved in 0.5% methylcellulose (Sigma-Aldrich) to create a 6% (w/v) solution, which was autoclaved prior to administration. Mice were gavaged with 150 μL of the carmine solution between 8:00am and 8:30am local time; animals were not fasted beforehand. Fecal pellets were collected every 30 minutes (up to 8 hours from time of gavage) and streaked across a sterile white napkin to assay for the presence of the red carmine dye. The time from gavage to initial appearance of carmine in the feces was recorded as the total intestinal transit time for that animal (*38*).

#### Restraint stress test

The reactivity of the HPA axis to moderate stress was determined using a restraint stress test (SRT). Each mouse was placed in a 50 ml plastic tube (11.4 x 2.8 cm) with holes for ventilation and an aperture in the cap for the tail. Blood samples (approx. 50μl) are collected from the tail vein into EDTA tubes (Fisher Scientific, USA) immediately after mice are placed in the tube (0 minutes), after 15 minutes spent in the tube, and 75 min after removal from the tube. Cellular constituents are separated by centrifugation (5 min, 14,800g) by mini-centrifuge. Plasma was stored at −80 °C until analysis of corticosterone concentration (Cayman chemical, USA). The SRT was exclusively performed between 09:00h and 11:00h, i.e., in the trough of the circadian rhythm of glucocorticoid secretion (*37*).

#### Cardiac perfusion and brain dissection

Animals that underwent MRI were sacrificed immediately after scanning. Each animal was deeply anesthetized with the appropriate isoflurane inhalation during MRI scanning and the subsequent terminal surgery. Non-MRI animals were also deeply anaesthetized with the appropriate isoflurane inhalation. Further, we confirmed each animal was in a deep level of anesthesia by pressing the toe with forceps and animal must be unresponsive. After confirmation that animals were in an appropriate level of anesthesia, we performed cardiac perfusion using 0.9% saline solution. We euthanized the animals immediately upon completion of cardiac perfusion via cervical dislocation. Immediately following cardiac perfusion, brains were removed from the skull and the two hemispheres were separated. In 50% of animals, the right hemisphere was placed in a 15 ml Falcon centrifuge tube partially filled with SuperGolgi Kit Solution A (Bioenno Tech, LLC, California, USA) for the analysis of spine morphology and density (*6*, *82*); see following section for further details. In the other 50% of animals, we used a Zivic mouse brain slicer tool, scalpel blades, and 1.5 mm and 1.2 mm tissue punches to collect samples of right prefrontal cortex (anterior cingulate and prelimbic area), nucleus accumbens, amygdala, and ventral hippocampus. Each region was stored in 4% paraformaldehyde for electron microscopy of axonal phenotypes. Samples of the left prefrontal cortex, nucleus accumbens, amygdala, and ventral hippocampus were also collected and placed in RNA later (Invitrogen-Thermo Fisher) for future gene expression studies.

### Dendritic morphology and spine density: Golgi-Cox staining

Golgi-Cox staining was performed by using the Super Golgi-Cox kit (Bioenno Tech, LLC, California, USA), according to the manufacturer’s instructions. The right hemisphere was impregnated with SuperGolgi Kit solution A for 10 days with the solution renewed after 2 days of immersion. After completion of impregnation, we rinsed the tissue blocks with deionized water (dH2O), then transferred them into post-impregnation buffer solution B and stored them at room temperature in the dark. We renewed the solution after one day of immersion. After 2 days of post-impregnation tissues were ready for sectioning. We performed coronal sectioning of 150 μm thickness by using a vibratome. Slices were collected into mounting buffer and mounted on slides. After washing the slides with 0.01 M PBS-T for 20∼30 min, we placed the slides into a closed staining jar with staining solution C in a dark area for 20 min. After that, slides remained in post-staining buffer for 20 min. We then performed thionin staining after PBS-T washing for proper visualization of structure. Further, slices were dehydrated with ethanol and xylene and cover slipped with DPX mounting medium, and slides were coded with various experimental groups.

### Analysis of dendritic morphology and spine density

Dendritic length and volume in hippocampus, amygdala, nucleus accumbens, and prefrontal cortex, were examined with microscopy (GE DeltaVision) under 10X magnification while spine density was calculated using images taken under 40X magnification. Images of pyramidal neurons in prefrontal cortex, hippocampus, and amygdala, and medium spiny neurons in nucleus accumbens were captured and analyzed using Imaris software. Dendritic length and volume measurements of individual neurons in each mouse were measured using the manual tracing option on Imaris and were subsequently averaged to obtain average dendritic length and average dendritic volume data per mouse.

For each animal, 2-11 neurons were analyzed for dendritic length and volume (in PFC, HUM1: 41 pyramidal neurons, HUM2: 16 pyramidal neurons, SPF: 23 pyramidal neurons, GF: 18 pyramidal neurons; in nucleus accumbens, HUM1: 41 medium spiny neurons, HUM2: 23 medium spiny neurons, SPF: 18 medium spiny neurons, GF: 24 medium spiny neurons; in amygdala, HUM1: 29 pyramidal neurons, HUM2: 26 pyramidal neurons, SPF: 22 pyramidal neurons, GF: 18 pyramidal neurons; in hippocampus, HUM1: 22 pyramidal, HUM2: 17 pyramidal neurons, SPF: 11 pyramidal neurons, GF: 18 pyramidal neurons.

For each animal, 2-6 neurons were analyzed for spine density (in PFC, HUM1: 41 pyramidal neurons, HUM2: 18 pyramidal neurons, SPF: 23 pyramidal neurons, GF: 17 pyramidal neurons; in nucleus accumbens, HUM1: 42 medium spiny neurons, HUM2:24 medium spiny neurons, SPF:15 medium spiny neurons, GF: 24 medium spiny neurons; in amygdala, HUM1:25 pyramidal neurons, HUM2: 21 pyramidal neurons, SPF:24 pyramidal neurons, GF:13 pyramidal neurons; in hippocampus, HUM1: 14 pyramidal neurons, HUM2: 14 pyramidal neurons, SPF: 3 pyramidal neurons, GF: 9 pyramidal neurons.

### Axon Myelination-Transmission Electron Microscopy

After primary fixation, samples were washed with 0.1M phosphate buffer and postfixed with 1% osmium tetroxide in 0.1M phosphate buffer and further dehydrated in a gradient series of acetone and samples were embedded in Spurr resin. For each specimen 70 nm thin sections were obtained from polymerized block using a Power Tome Ultramicrotome (RMC, Boeckeler Instruments. Tucson, AZ). The thin sections were stained with uranyl acetate and lead citrate stain. Further, images were examined by using a JEOL 1400 Flash Transmission Electron Microscope (Japan Electron Optics Laboratory, Japan) at an accelerating voltage of 100kV. Electron micrographs were obtained from the different areas of brain including amygdala, prefrontal cortex, hippocampus, and nucleus accumbens. Each axon was labelled based on decompaction level and received a label of “none,” “low,” “high,” or “partial.” While these labels were subjective, they were all performed by the same rater to ensure consistency. Examples of each type of label are shown in Figure 9(a). While axon diameter was measured regardless of decompaction level, myelin sheath thickness was only measured in axons with no or low decompaction. Axons with high decompaction were not included in the laminal count analysis.

**Figure 9.**
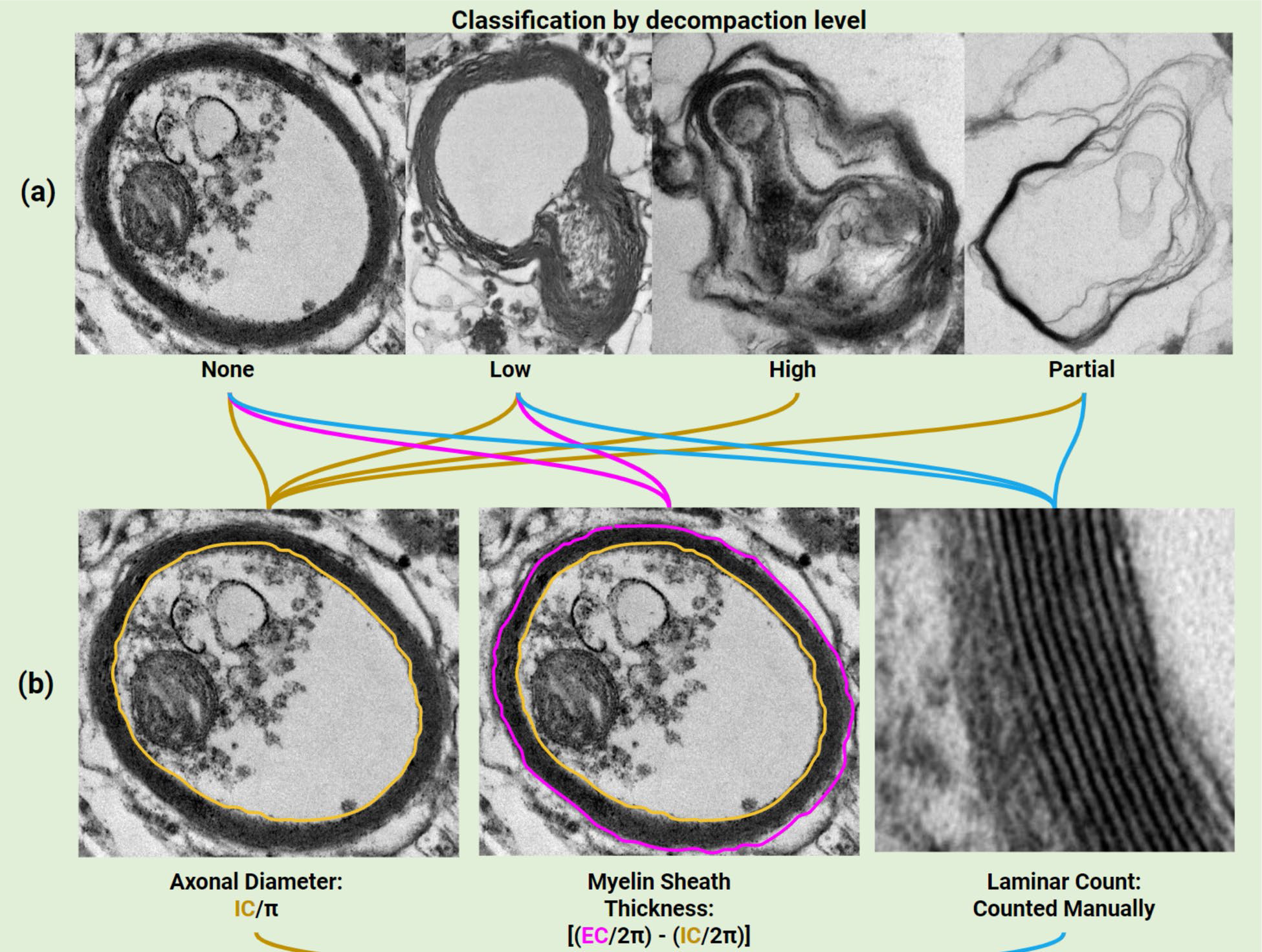
Electron microscope analysis of axons. (a) a single rater classified the axons by decompaction levels (none, low, high, and partial), example images of each level are shown. Axonal diameter was calculated for all classes; only axons classified as none or low were assessed for myeline sheath thickness; axons classified as highly decompacted were not included in laminar counts, (b) methods for generating axonal measurements. IC is internal circumference and EC is external circumference.

Axon diameter was measured using ImageJ. The line selection tool was used, and a freehand line traced the interior of the axon, which was then measured using the measure tool and taken to be the inner circumference (IC) of the axon. The measurement had already been calibrated to the actual scale of the axon using the scale bar in each image and the Set Scale Function in ImageJ. The inner circumference was then divided by π to calculate the internal diameter of the axon. Similarly, myelin sheath thickness was measured with the line selection tool. A freehand line traced the exterior of the axon and represented the external circumference (EC) of the axon. IC and EC were put into the following equation to calculate myelin sheath thickness: [(EC/2π) - (IC/2π)]. Further, individual layers of myelin, or laminae were manually counted at four different points in the axon from images with high magnification (See Figure 9(b)).

For axonal diameter analysis, 1-44 axons were analyzed from each animal (in PFC, HUM1: 109, HUM2: 78, SPF=35, GF=75; in nucleus accumbens, HUM1: 113, HUM2: 133, SPF=60, GF=70; in amygdala, HUM1: 139, HUM2: 102, SPF=87, GF=116; in hippocampus, HUM1: 166, HUM2: 126, SPF=87, GF=113).

For myelin sheath analysis, 1-34 axons were analyzed from each animal (in PFC, HUM1: 22, HUM2: 23, SPF=21, GF=36; in nucleus accumbens, HUM1: 30, HUM2: 71, SPF=12, GF=38; in amygdala, HUM1: 87, HUM2: 55, SPF=51, GF=62; in hippocampus, HUM1: 79, HUM2: 46, SPF=51, GF=62).

For laminar count analysis, from each animal, 1-32 axons were analyzed (in PFC, HUM1: 90, HUM2: 40, SPF=0, GF=55; in nucleus accumbens, HUM1: 68, HUM2: 60, SPF=0, GF=51; in amygdala, HUM1: 120, HUM2: 61, SPF=0, GF=84; in hippocampus, HUM1: 98, HUM2: 74, SPF=0, GF=97). SPF animals were not included in this analyses due to high levels of decompaction which prevented accurate laminar counts.

### Gut microbiota analyses of inocula and mouse cecal contents

#### Extraction of microbial DNA

To determine engraftment of the donor profile, we collected post-mortem cecal contents from adult offspring between 10:00 am to 7:00 pm. (Figure 7). The time from inoculation to collection of cecal content varied from 11 to 13 weeks. Samples were collected in 1.5ml Eppendorf tubes and immediately stored on dry ice in a −80 °C freezer. Genomic DNA was extracted from cecal content samples and the inocula using the FastDNA Spin kit for soil (MP Biomedicals, USA), according to the manufacturer’s instructions. DNA concentration and purity were measured with a Qubit Flex Fluorometer (Invitrogen, Thermo Fischer Scientific, USA). Library preparation and 16S rRNA gene amplicon analysis were performed by the Michigan State University Genomics core.

#### 16S rRNA amplicon sequencing

The V3-V4 hypervariable regions of the 16S rRNA gene were amplified using single barcoded, Illumina-compatible primers following the protocol described in Caporaso, JG, et al. (2011) (*118*), except that the 515f universal primer described in the paper was replaced with the 341f universal primer, upstream of the V3 region. PCR products were batch normalized using an Invitrogen SequalPrep DNA Normalization plate, and the product recovered from the plates pooled. The pools were cleaned up and concentrated using a QIAquick Spin column and AMPure XP magnetic beads. The pool was QC’d and quantified using a combination of Qubit dsDNA HS, Agilent 4200 TapeStation HS DNA1000, and Invitrogen Collibri Illumina Library Quantification qPCR assays. This pool was loaded onto the MiSeq v2 Standard flow cell and sequencing was carried out in a 2×250bp paired-end format using a MiSeq v2 500 cycle reagent cartridge. Custom sequencing and index primers complementary to the 341f/806r oligomers were added to appropriate wells of the reagent cartridge. Base calling was done by Illumina Real Time Analysis (RTA) v1.18.54, and the output of RTA was demultiplexed and converted to FastQ format with Illumina Bcl2fastq v2.20.0.

#### Bioinformatics of microbiome data

Raw FASTQ files were processed using QIIME2. We imported single-end fastq (forward reads) sequence files to QIIME2 and performed demultiplexing using the q2-demux plugin. After demultiplexing and quality filtering, the total reads per sample ranged from 141823 to 414669. One sample was excluded due to very low reads. The sequences were then trimmed to remove primers (cutadapt plugin) and chimeric reads using DADA2 plugin. Reads with quality score Q>20 were kept, and the read length was 250bp. Phylogenetic trees were generated using the align-to-tree-mafft-fasttree command in QIIME2. Taxa were assigned based on the SILVA 16S rRNA database (version 138). Diversity analysis was done with a rarefaction set at 103790, which retained 4,977,600 (60.57%) features. Alpha and beta diversity analyses were conducted using a diversity plugin (*119*).

## Statistical analysis

Statistical analysis was performed with R version 4.2.0. Statistical significance was determined using three-way ANOVA or, for longitudinal data, mixed model ANOVA with repeated measures followed by post hoc Tukey’s test or –t-test. We used two-way ANOVA followed by post hoc Tukey’s test for sex-stratified analyses.. The effective number of independent tests (Meff) was calculated using Li and Ji (2005) method (*120*). Correction for multiple comparisons was done using the Bonferroni method. For behavioral data analysis, Meff was 17 and p-value <0.003 was considered statistically significant. For dendritic morphology, Meff was 7 and p-value <0.007 was considered statistically significant. For myelination data analysis, Meff was 11 and p-value <0.0045 was considered statistically significant. For sucrose preference, stress reactivity, and red carmine test, a p-value of <0.05 was considered statistically significant.

## Acknowledgements

This study was supported by NIH Grants P30 DK34987 to M.A.A. and R33 MH104330 to R.C.K. and through startup funds provided by Michigan State University to R.C.K.

Author contributions are as follows: Conceptualization: R.C.K, Methodology: R.C.K, C.B., S.L., A.C., M.A.A., L.M., Software: A.A., Statistical analysis: A.A. and M.A.A., Investigation: H.D., R.R., C.B., S.L., A.W., A.C., M.A.A., Resources: R.C.K., Data Curation: H.D., A.A., A.C., Writing – Original Draft: H.D., Writing – Review and Editing: All authors, Visualization: R.C.K., H.D., R.R., A.A., C.B. M.A.A., Supervision: R.C.K., Project administration: H.D., Funding acquisition: R.C.K and M.A.A.

We thank the staff at the Michigan State University RTSF Genomics Core, especially Kevin Childs and Emily Crisovan, who generated the 16S rRNA sequencing data for the inocula and mouse cecal samples. We thank the staff at Michigan State University’s Center for Advanced Microscopy, especially Alicia Withrow, for generating the electron microscopy images. We thank the staff at Michigan State University’s Investigative Histopathology Laboratory, especially Amy Porter and JessieLee Neumann, for their assistance with the Golgi-Cox staining. We thank the staff at the UNC Microbiome Core for carrying out 16S rRNA sequencing and bioinformatic analysis of human donor samples. We thank Jennifer Quesenberry, Wendy Neuheimer, Dianne Evans, and Preman Koshar who served as study coordinators for the human infant study that provided the donor samples for this work. Corey Shope and DeSiyre Spurgeon provided additional administrative assistance for that study. We thank the families for their participation.

The authors declare that they have no competing interests.

All group-level data needed to evaluate the conclusions in the paper are present in the paper and/or the Supplementary Materials. Individual level data can be provided by Dr. Knickmeyer and may require scientific review and a completed data use or material transfer agreement. Requests for individual-level data should be submitted to.

