## Supplementary Table S1 for "Effect of Human Infant Gut Microbiota on Mouse Behavior, Dendritic Complexity, and Myelination"

Harikesh Dubey *et al.*

**This PDF file includes:**

Table S1

**Table S1.**

Checklist for Guidelines for Reporting on Animal Faecal Transplantation (GRAFT)

### GRAFT FRAMEWORK

#### Guidelines for Reporting on Animal Faecal Transplantation

CONSISTENCY . TRANSPARENCY . TRANSLATION

GRAFT

#### COLLECTION

| | | Section / Paragraph |
|---|---|---|
| **Donor phenotype / characteristics** | a. Number of individual donors (per group)<br>b. Detailed description of donor characteristics (see also ARRIVE Guidelines for animal donors), including but not limited to:<br>- Species / strain of donors<br>- Sex / gender of donors<br>- Age and developmental stage of donors<br>c. Details of control and experimental phenotypes (e.g. healthy vs disease phenotype)<br>- Inclusion and exclusion criteria, with particular attention to factors relevant to the microbiome (e.g. diet, exercise)<br>d. Details on housing and husbandry<br>- Facility specifications (i.e. SPF / GF; If GF, include specifications of animal unit / isolator)<br>- Co-housing vs single-housing<br>- Arrangement of cages across racks (particularly with regards to separation of donor groups and separation from FMT recipient animals)<br>- Bedding and chow | a. Section-Donor phenotype/characteristics /paragraph-1<br>b. Section-Donor phenotype/paragraph-last<br>c. Section-Donor phenotype/characteristics /paragraph-last<br>d. Section-Donor phenotype/characteristics /paragraph-last, and section-Recipient Animals and Fecal Transplant Procedure/ paragraph-1 |
| **Sample collection process** | a. Type of sample (i.e. faecal pellet, intestinal / caecal content, mucosal scraping)<br>b. Time of day of collection and details on minimisation of circadian rhythm effects<br>c. Animal handling during collection<br>d. Details on sample collection methods (e.g. placing animal into clean cage until defecation or direct post-mortem collection from caecum or intestines)<br>- HUMAN DONORS: collection methods (e.g. normal defecation or directly from specific region of intestines during colonoscopy, medically-indicated or otherwise) | a. Section-Donor Sample Collection and Storage/paragraph-1 and last<br>b. Section-Donor Sample Collection and Storage/paragraph-1 and |
| **Measures to minimise contamination** | a. Aseptic procedures and protocols adopted during and after collection | a. Section-Donor Sample Collection and |
| **Measures to maximise viability** | a. Time from sample collection until anaerobic conditions / freezing / administration<br>b. Details of anaerobic conditions during collection / transport | a. Section-Creation and storage of inoculums<br>[illegible] |
| **Immediate storage conditions** | a. Methods to minimise oxidative stress (i.e. use of transport medium)<br>b. Immediate storage conditions (e.g. stored in reduced medium, snap frozen in liquid nitrogen, kept on ice or at ambient temperature etc.)<br>c. Details on pooling of samples (if relevant)<br>- Method of pooling (e.g. equal weight of initial sample from each donor prior to processing or equal volume of processed liquid)<br>- Number of individual donors within each pool | a. Section-Donor Sample Collection and Storage/paragraph-1<br>b. Section-Donor Sample Collection and Storage/paragraph-1 and last |

#### PROCESSING

| | | Section / Paragraph |
|---|---|---|
| **Vehicle preparation** | a. Details of solution, including formulation, concentration, pH, temperature, volume<br>b. Additives used to support microbial viability<br>c. If de-oxygenated solution is used, specify method of de-oxygenation | a. Section-Donor Sample Collection and Storage and section- Creation and |
| **Concentration** | a. Report using standardised units (mg/ml)<br>- Avoid inaccurate units (e.g. pellets/ml) | a. Section- Creation and storage of inoculums |
| **Homogenisation method** | a. Equipment used (e.g. vortex, Stomacher, autoclaved spatula)<br>b. Intensity (using standardised units where possible)<br>c. Time and temperature | a. Section- Creation and storage of inoculums<br>b. Section- Creation and |

### GRAFT FRAMEWORK

#### Guidelines for Reporting on Animal Faecal Transplantation

CONSISTENCY . TRANSPARENCY . TRANSLATION

GRAFT

#### PROCESSING cont.

| | | Section / Paragraph |
|---|---|---|
| **Filtration method** | a. Method of filtration (e.g. gravity, centrifuge, strainer, stomacher bag)<br>- Centrifuge: specify time, xg and temperature<br>- Gravity: specify time and conditions (i.e. ambient, anaerobic, temperature)<br>- Physical strainer / membrane: specify pore size or equivalent detail and filtration method | None |
| **Anaerobic conditions** | a. Clearly state if / when anaerobic conditions were used<br>b. Details of anaerobic conditions (i.e. chamber type, gas mix, temperature etc.) | a. Section- Creation and storage of inoculums |
| **Quality control** | a. Method used to assess FMT quality and composition prior to administration (e.g. plating, genomic sequencing)<br>b. Outcome of quality assessment (e.g. CFU/ml, diversity index) | a. Section- Creation and storage of inoculums, 16 s rRNA (conducted after |

#### STORAGE

| | | Section / Paragraph |
|---|---|---|
| **State of final product** | a. Define administered product as:<br>- Faecal slurry (i.e. faecal contents with minimal filtration) - or -<br>- Faecal supernatant / filtrate (i.e. microbial free) - or -<br>- Microbial preparation (i.e. lyophilised or other) | a. Section- Creation and storage of inoculums |
| **Time in storage** | a. Time between preparation of final product and administration | a. Section- Creation and [illegible] |
| **Storage conditions** | a. Details of storage conditions between preparation and administration, including:<br>- Volume per aliquot<br>- Storage temperature<br>- Duration of storage<br>b. If faecal product is used fresh, this must be clearly stated with details including:<br>- Short term storage conditions (i.e. on ice, fridge, room temperature, anaerobic chamber)<br>- Time between preparation and administration | a. Section- Creation and storage of inoculums<br>b. Section- Creation and storage of inoculums |
| **Freeze / thaw cycles** | a. Method of thawing faecal product prior to administration<br>- Include number of freeze-thaw cycles | a. Section- Creation and storage of inoculums |

#### RECIPIENT PREPARATION

| | | Section / Paragraph |
|---|---|---|
| **Recipient phenotype / characteristics** | a. Number of recipient animals (per group)<br>- If multiple animals receive FMT from the same donor (or pooled sample), this number should be reported for each donor, separately to the total<br>b. Detailed description of recipient characteristics (see also ARRIVE Guidelines), including but not limited to:<br>- Species / strain of recipients<br>- Sex of recipients<br>- Age and developmental stage of recipients<br>c. Details on housing and husbandry<br>- Facility specifications (i.e. SPF / GF; If GF, specifications of animal unit / isolator)<br>- Co-housing vs single-housing<br>- Arrangement of cages across racks (particularly with regards to separation of experimental groups and separation from FMT donor animals)<br>- Bedding and chow | a. Section- Recipient Animals and Fecal Transplant Procedure/paragraph-1<br>b. Section- Recipient Animals and Fecal Transplant Procedure/paragraph-1<br>c. Section- Recipient Animals and Fecal Transplant Procedure/paragraph-1 and section- Housing Conditions at Michigan [illegible] |

### GRAFT FRAMEWORK

#### Guidelines for Reporting on Animal Faecal Transplantation

CONSISTENCY . TRANSPARENCY . TRANSLATION

GRAFT

#### RECIPIENT PREPARATION cont.

| | | Section / Paragraph |
|---|---|---|
| **Host preparation techniques** | a. Methods of host preparation used prior to transplantation (e.g. antibiotic depletion, bowel cleansing, fasting) with relevant detail, including but not limited to:<br>- Duration<br>- Frequency (e.g. of changing antibiotic solution)<br>- Specific treatment used (e.g. antibiotic names and concentrations)<br>b. Preparation methods used in control group(s), with details as above<br>c. Adverse events in response to preparation treatment (e.g. weight loss with antibiotics) | a. Section- Recipient Animals and Fecal Transplant Procedure/paragraph-2<br>b. Section- Recipient Animals and Fecal Transplant |
| **Confirmation of preparation success** | a. Ideally, successful depletion of recipient microbiota should be confirmed through faecal analysis prior to FMT | a. Section- Recipient Animals and Fecal [illegible] |

#### ADMINISTRATION

| | | Section / Paragraph |
|---|---|---|
| **Route and method of administration** | a. Oral or rectal administration (or both)<br>b. Method of administration (e.g. oral gavage, lavage, enema, coprophagia)<br>c. Details on use of anaesthesia or fasting prior to administration (particularly rectal) and coprophagic approaches (i.e. was additional FMT smeared on coat to improve uptake) | a. Section- Recipient Animals and Fecal Transplant Procedure/paragraph-2 |
| **Volume and concentration** | a. Define in standard units for each individual FMT<br>- Specify if absolute unit or relative to body weight of recipient | a. Section- Recipient Animals and Fecal |
| **Time and frequency** | a. Time of day of administration<br>b. Frequency of FMT, including total number and daily frequency (i.e. a total of 3 FMT by oral gavage at a frequency of 1 per day, number of days between doses)<br>c. Time between FMT administration and assessment of outcomes (i.e. disease status, behavioral change, microbiota composition etc.) | a. Section- Recipient Animals and Fecal Transplant Procedure/paragraph-2<br>b. Section- Recipient |
| **Control treatment** | a. Define treatment received by control animals (e.g. vehicle solution, autologous transplant, heat-killed FMT, FMT from control donor group)<br>- Include control formulation, concentration, volume, time, and frequency as above | a. Section- Recipient Animals and Fecal Transplant |

#### CONFIRMATION

| | | Section / Paragraph |
|---|---|---|
| **Engrafting / uptake of donor profile** | a. Define how engraftment / uptake of the FMT procedure was confirmed (e.g. 16S rRNA / shotgun sequencing, faecal culture)<br>- It is recommended that the same analysis be applied to the final FMT product administered to compare composition of donor and recipient<br>b. Timing of sample collection for engraftment assessment relative to FMT administration and outcome assessments<br>c. Details on sample collection methods, as for donor:<br>- Time of day of collection<br>- Handling during collection<br>- Method: Placing animal into clean cage until defecation or direct post-mortem collection from colon, caecum or other site | a. Section- Gut microbiota analyses/sub section-Extraction of microbial DNA Subsection-16s amplicon sequencing and metagenomics analysis<br>b. For timeline Fig 7, Section-Gut microbiota analyses/sub section-Extraction of |
| **Durability / stability of donor profile** | a. Particularly for lengthy experimental designs, it may be informative to analyse the recipient microbiota at multiple time-points after FMT administration to determine the long-term stability of the donor profile within the recipient | a. Section-Gut microbiota analyses/sub section-Extraction of |

**Other Supplementary Materials for this manuscript include the following:**

Data S1 to S7

**Data S1. (Separate file)**

**Microbial taxonomic composition of cecal contents varies between humanized mice and specific-pathogen free and germ-free controls.** **(a)** Comparison of alpha diversity across offspring cecal contents using Shannon index, **(b)** Comparison of alpha diversity across offspring cecal contents using Chao1 index, **(c)** Alpha diversity differences in original human inocula (HUM1 and HUM2) and mouse SPF inoculum, **(d)** Comparison of β-diversity across offspring cecal contents at the genus level using principal coordinate analysis (PCoA) and PERMANOVA based on unweighted unifrac method, **(e)** Comparison of β-diversity across offspring cecal contents at the genus level using principal coordinate analysis (PCoA) and PERMANOVA based on weighted unifrac method, **(f)** Comparison of β-diversity across offspring cecal contents at the genus level using principal coordinate analysis (PCoA) and PERMANOVA based on Bray-Curtis method, **(g)** Comparison of β-diversity across offspring cecal contents at the genus level using principal coordinate analysis (PCoA) and PERMANOVA based on Jaccard Index, **(h)** Variation in relative abundance of microbial genera per sample visualized using a taxa barplot, **(i)** Variation in relative abundance of microbial genera in HUM1, HUM2, and SPF inocula using a taxa barplot **(j)**, Summary of the establishment of core donor amplicon sequence variants (ASVs) into recipient mice for HUM1, HUM2 and SPF inocula.

**Data S2. (Separate file)**

**Microbiome inoculation group influenced anxiety-related and exploratory/locomotor behaviors, but not social behavior or recognition memory.** Statistical models and violin plots comparing HUM1, HUM2, SPF, and GF mice on **(a)** elevated plus maze, **(b)** light/dark test, **(c)** open field test, **(d)** novel object recognition test, and **(e)** three-chamber social test.

**Data S3. (Separate file)**

**Microbiome inoculation group influenced depressive behavior and stress responses.** Statistical models and associated figures comparing HUM1, HUM2, SPF, and GF mice. **(a)** sucrose preference test, **(b)** repeated measures analysis of hormonal responsivity, indexed by corticosterone levels, to a restraint stress test, **(c)** secondary analyses looking at microbiome effects on change in corticosterone from baseline to 15 minutes (animal still in restraint tube), change from 15 minutes to 90 minutes (animal no longer in tube), and change from baseline to 90 minutes

**Data S4. (Separate file)**

**Minimal effects of microbiome inoculation group on intestinal transit time using the red carmine test.**

**Data S5. (Separate file)**

**Gut microbiome may influence myelin sheath thickness in the hippocampus, but has minimal effects on other regions and on axonal diameter, laminar counts, and g-ratio.** Statistical models and violin plots comparing axonal phenotypes in different regions of HUM1, HUM2, SPF, and GF mice brains. Data derived from electron microscopy. **(a)** Myelin sheath thickness **(b)**, axon diameter **(c)**, laminar count **(d)**, g-ratio.

**Data S6. (Separate file)**

**Dendritic morphology differs between gut microbiome inoculation groups.** Statistical models and violin plots comparing HUM1, HUM2, SPF, and GF mice for dendritic phenotypes assessed via Golgi-Cox staining in different brain regions. **(a)** dendritic length, **(b)** dendritic volume, **(c)**, dendritic spine density**.**

**Data S7. (Separate file) Animal details.** This file provides a list of all offspring animals in the described experiments with their microbiome group assignment, cohort, gender, and whether they were included in the MRI, Golgi-Cox, and Electron Microscopy components of the the project.
